# A Chromosome-scale Genome Assembly of *Meloidogyne hapla* Reveals Localized Recombination Hotspots Enriched with Effector Proteins

**DOI:** 10.1101/2025.05.21.654606

**Authors:** Pallavi Shakya, Muhammad I. Maulana, Etienne GJ Danchin, M. Laurens Voogt, Stefan J.S. van de Ruitenbeek, Jacinta Gimeno, Adam P Taranto, Alison C. Blundell, Evelin Despot-Slade, Nevenka Meštrović, Ana Zotta Mota, Dadong Dai, Valerie M. Williamson, Mark G. Sterken, Shahid Siddique

## Abstract

Root-knot nematodes (*Meloidogyne spp.)* are among the most destructive agricultural pests that cause significant yield losses across a wide range of crops. *Meloidogyne hapla,* a diploid species, is a valuable model for studying root-knot nematodes due to its parasitic diversity, small genome, and a reproductive strategy that facilitates genetic analysis. Here, we present a high-quality chromosome-scale assembly of *M. hapla,* generated using multiple sequencing platforms–PacBio HiFi, ONT, Illumina and HiC. The 59 Mb assembly comprises 16 chromosome-length scaffolds, notably lacking canonical telomeric repeats. Instead, we identified a tandem 16-mer repeat mainly present at scaffold ends, suggesting an alternative system for chromosome-end maintenance. Genetic linkage analysis of F2 populations derived from crosses between *M. hapla* strains validated the assembly but also revealed anomalies indicating chromosome structure differences between parental isolates such as fissions, fusions, and rearrangements. This analysis also revealed sharply delineated zones of high recombination on most chromosome arms. We also identified 1,258 genes encoding putative secreted proteins (PSP), which should be enriched in genes involved in host interaction and pathogenicity. Most of the PSP genes had orthologs in other plant parasitic nematode species, and the majority were pioneers, lacking known functional domains. Notably, we found that PSPs are significantly enriched in high-recombination zones, possibly facilitating their rapid evolution. Overall, our study provides new insights into the genome structure of diploid root-knot nematodes and highlights the interplay between genome architecture, recombination, and parasitism. These findings raise new questions about how genetic and genomic adaptations drive the success of root-knot nematodes as plant parasites.

## Introduction

Nematodes are among the most abundant and diverse animal phyla, containing an estimated 1-10 million species (Hugot et al., 2001). This remarkable diversity is reflected in their ecological niches—ranging from free-living forms in soil and water to parasitic species infecting plants and animals. Among the approximately 4,100 species identified as plant parasites, species in the genus *Meloidogyne* (commonly known as root-knot nematodes, or RKNs) stand out as the most economically damaging (Trudgill & Blok, 2001; Bird et al., 2009). The four most damaging species, *M. arenaria*, *M. incognita*, *M. javanica*, and *M. hapla,* possess an extraordinary host range spanning a diverse array of crop species (J. T. Jones et al., 2013). Three of these species (*M. arenaria*, *M. incognita*, *M. javanica*) are closely related (*Meloidogyne* Clade I) and are globally distributed in tropical and subtropical regions. In contrast, *Meloidogyne hapla* (Clade II) is generally found in more temperate climates and parasitizes a broad range of host plants, although isolates differ in their host range, pathogenicity, and behavior (Beek et al., 1998; Liu & Williamson, 2006; Melakeberhan et al., 2012; Mitkowski & Abawi, 2003).

The RKN lifecycle generally spans about a month and encompasses six distinct stages consisting of embryo, four juvenile stages (J1-J4) and an adult stage (Escobar et al., 2015; Páez, 2023). As obligate sedentary endoparasites, RKNs spend most of their life cycle feeding from a permanent site within the root vascular system. These feeding sites are characterized by nematode-induced multinucleated giant cells that act as nutrient sinks for the nematodes, and by distinctive root galls or “knots”, a hallmark of RKN infestation (Lin & Siddique, 2024; Rutter et al., 2022). How RKNs establish these specialized feeding sites in such a broad range of plant species, spanning monocots, dicots, annuals, and perennials is a question of both scientific and practical interest. Nematode genes responsible for establishing feeding sites or contributing to differences in host range are largely unknown. However, several proteins secreted by plant-parasitic nematodes have been demonstrated to contribute to parasitism and are commonly referred to as effectors (Bali & Gleason, 2024; Molloy et al., 2024; Rehman et al., 2016; Siddique et al., 2022). These effectors include genes likely acquired via horizontal transfers as well as many encoding pioneer proteins with no known protein motifs.

*Meloidogyne* spp. exhibits a wide range of karyotypes and reproductive mechanisms (Triantaphyllou & Hirschmann, 1980). Whereas most Clade I species reproduce asexually without meiosis and carry genomes with various degrees of polyploidy, most isolates of *M. hapla* are diploid and reproduce by facultative meiotic parthenogenesis (Beek et al., 1998; Triantaphyllou, 1966). Sexual reproduction occurs when females are fertilized by migratory males, and sperm-oocyte fusion occurs to generate hybrid offspring (Triantaphyllou, 1966; Liu et al., 2007a). However, cytological studies have shown that, in the absence of males, sister chromatids of meiosis II are rejoined to restore diploidy asexually (Triantaphyllou, 1966; Liu et al., 2007b). This reproductive mechanism has facilitated controlled crosses between strains and the production of F2 lines. Molecular marker analysis of F2 lines from a cross of two different strains showed these lines are largely homozygous for sequence polymorphisms and thus resemble recombinant inbred lines (RILs) (Liu et al., 2007a). Additionally, these F2 lines have been used to produce a marker-based genetic map and to identify genetic loci affecting interactions with host and/or behavior (Wang et al., 2010; Thomas et al., 2012; Thomas & Williamson, 2013; Guo et al., 2017).

Our study focuses on *M. hapla* for its genetic tractability and diploid genome, which make it a favorable model and therefore reference organism for RKN genetics and genomics. A previous draft genome sequence of the inbred *M. hapla* strain VW9 has been produced, and a DNA-based linkage map was generated based on segregation of polymorphisms in RIL-like F2 lines from a cross between strain VW9 and another nematode strain (VW8) (Opperman et al., 2008). However, this genome assembly is highly fragmented, limiting both the localization of genes responsible for phenotypic traits and synteny analyses with other RKNs. Recent advances in sequencing technologies have led to highly contiguous assemblies for several diploid and polyploid *Meloidogyne* species (Bali et al., 2021; Dai et al., 2023; Mota et al., 2023; Winter et al., 2024). In these studies, the canonical nematode telomeric repeats (TTAGGC)n were not detected at chromosome ends. Instead, in three Clade I species, *M. incognita*, *M. arenaria* and *M. javanica,* long arrays of species-specific complex tandem repeats were found, mostly enriched at one scaffold end (Dai et al., 2023; Mota et al., 2023). Furthermore, telomere associated proteins such as telomerase and shelterin complexes that are evolutionarily conserved in other nematode clades were not identified suggesting an alternative mechanism for chromosome end maintenance may be at play in RKN (Mota et al., 2023).

Here, we generated a *de novo* chromosome-level assembly of diploid *M. hapla* using complementary long-read sequencing strategies. We characterized the non-canonical repeat sequences at the end of chromosome-length scaffolds. Additionally, we utilized data from previously generated F2 lines to compare scaffold structure with genetic linkage groups. This analysis revealed a recombination landscape characterized by extraordinary recombination rates mostly on chromosome arms and provided evidence for differences in chromosome structure/behavior between isolates. To the best of our knowledge, this is the first chromosome-scale RKN genome assembly validated by genetic linkage analysis. Furthermore, we examine the chromosomal distribution of genes encoding secreted proteins (effector candidates) and provide evidence for their enrichment in the high recombination zones.

## Results

### Chromosome-level genome assembly of *Meloidogyne hapla* strain VW9

We produced a high-quality, chromosome-level genome assembly for *Meloidogyne hapla* strain VW9, the same strain used in a previous assembly by Opperman et al. (2008) (**PRJNA29083**). We used a combination of PacBio HiFi sequencing (56x coverage), Oxford Nanopore Technologies (ONT) long-read sequencing (143x coverage), Illumina short read sequencing (138x coverage), and Hi-C Chromatin conformation capture (263x coverage). Using Hifiasm with the HiFi, ONT, and Hi-C datasets, we produced an initial assembly with 36 contigs. This assembly was assessed for contaminants with Blobtools, where all 36 contigs were assigned the taxonomy Nematoda (**Supp. Fig. 1; Supp. Data 1).** This assembly was further polished with the Illumina reads and subsequently scaffolded using Juicer and 3D-DNA with the Hi-C data (**Fig. 1A**, **Supp. Data 1 & 2**). Additionally, the mitochondrial genome was assembled (**Supp. Fig. 2**). The resulting assembly consisted of 16 scaffolds for a total length of 59.2 Mb with an N50 of 3.8 Mb. The assembly size closely matched the 61.6 Mb genome size estimated by k-mer analysis (**Supp. Fig. 3A**) and was consistent with flow cytometry data, which estimated the diploid nuclear DNA content of *M. hapla* to be 121±3 Mb, corresponding to a haploid genome size of approximately 60 Mb (Blanc-Mathieu et al., 2017). Analysis of the Illumina data supported a diploid genome structure for *M. hapla* and almost entirely homozygous (**Supp. Fig. 3B)**. The high integrity of this assembly was further supported by the Merqury k-mer plot, which indicated that almost all the information present in Hi-Fi reads was captured in the haploid assembly (**Supp. Fig. 3C**). Overall, this new *M. hapla* genome assembly represents a significant improvement over the previous version and is the most contiguous so far for a RKN.

**Fig. 1:**
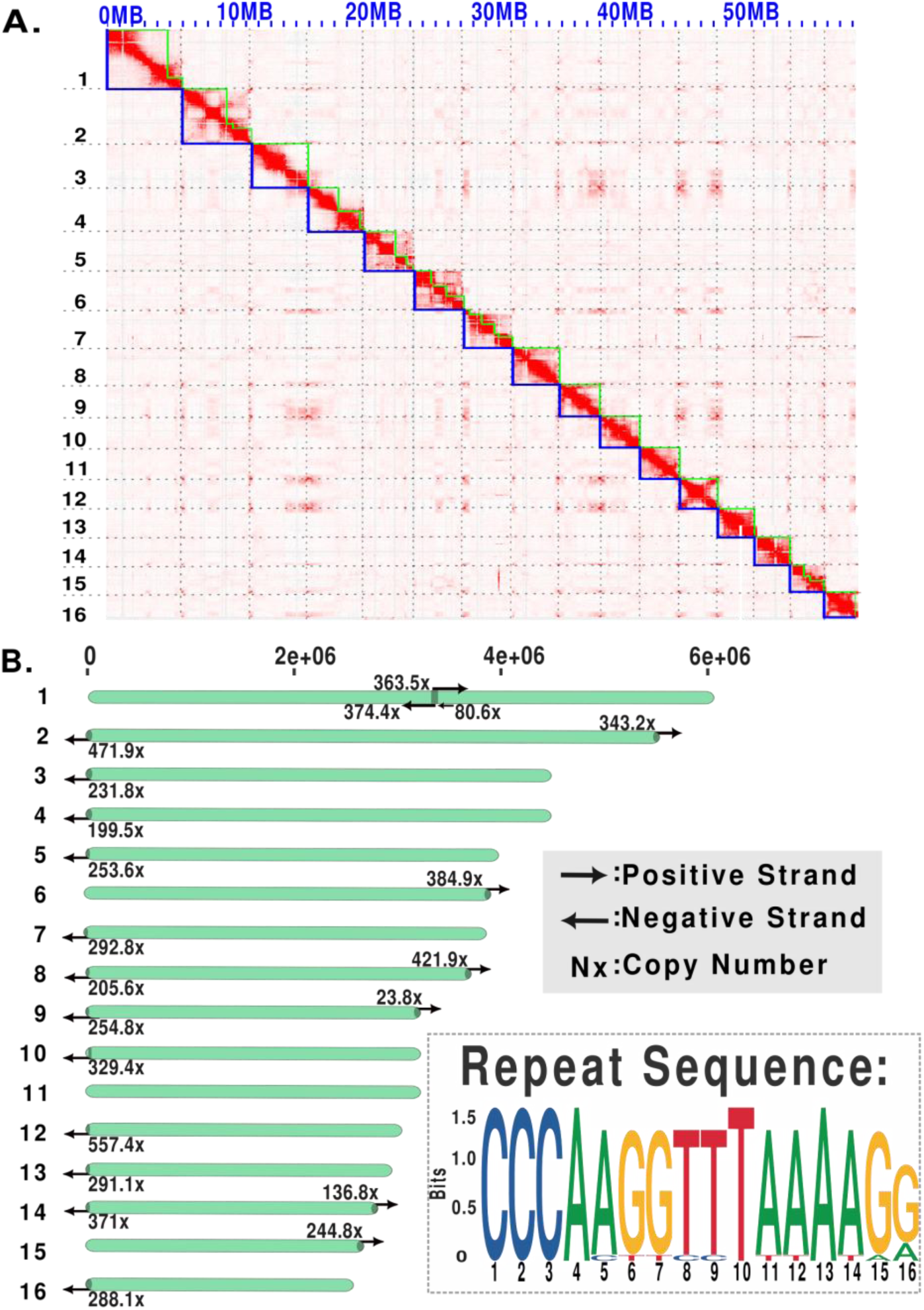
Genome Assembly and characteristics of *Meloidogyne hapla*strain VW9. **A.** HiC contact map of *M. hapla* showing 16 chromosome-scale scaffolds. Green lines denote the edges of contigs, and blue lines denote the edges of scaffolds. **B.** Distribution of the 16- mer repeats across chromosome-scale scaffolds. Numbers indicate repeat copy number, and arrows show repeat orientation within scaffolds

In the Hi-C contact map, we observed patterns suggesting physical proximity between scaffold tips of different chromosomes (**Fig. 1A**). To assess whether these patterns were an assembly artifact, we checked the quality of raw Hi-C reads and built Hi-C contact maps with different draft assemblies (**Supp. Fig. 4).** This analysis supports that the patterns are due to inter-chromosomal tip proximity.

To define the scaffold ends in the *M. hapla* assembly we analyzed the terminal 24 kb of each scaffold for tandem repeats and found arrays of a 16-mer with the consensus sequence (CCCAAGGTTTAAAAGG) at 17 scaffold ends. Nine scaffolds (S3, S4, S5, S6, S7, S12, S13, S15, S16) have this tandem repeat at only one end, while four scaffolds (S2, S8, S9, and S14) show the same tandem repeat at both ends (**Fig. 1B**). The length profile of these arrays ranges from approximately 1,289 bp, to 8,912 bp. For S10, a variant tandem repeat with consensus TTATAAAGGAAGTGGG is present starting 4 kb from one end. A search for these end-specific tandem repeat arrays within scaffolds identified only three arrays at the center of S1 scaffold (**Fig. 1B**; **Supp. Table 1)**. The internal arrays in S1 formed a large palindrome with over 300 copies of the repeat followed by over 300 copies of the complementary sequence. Among the scaffolds, only S11 entirely lacked either repeat array.

In addition, to validate whether the candidate telomeric repeats indeed localize to the chromosome ends, we performed fluorescence in situ hybridization (FISH) analysis on elongated chromosomes of *M. hapla* (**Fig. 2**) using the 16-mer repeat found at 17 scaffold ends. Due to the tiny size of the chromosomes and the difficult biological material, we were not able to obtain complete chromosome complement spreads. Nevertheless, for the intact chromosomes that were discernible, the 16-mer repeats predominantly localized at or near the termini of multiple chromosomes (**Fig. 2A, B**). Several chromosomes showed signal at only one end, while others displayed no signal (**Fig. 2B)**. The signal intensity varied between chromosomes, indicating differences in repeat array lengths. Furthermore, we detected a signal internally for at least one chromosome (**Fig. 2A**), corroborating the presence of an internal repeat array as observed in S1. Overall, FISH results confirm the 16-mer repeat is enriched predominantly at chromosome ends and most likely constitutes specific telomeric sequences in *M. hapla*.

**Fig. 2:**
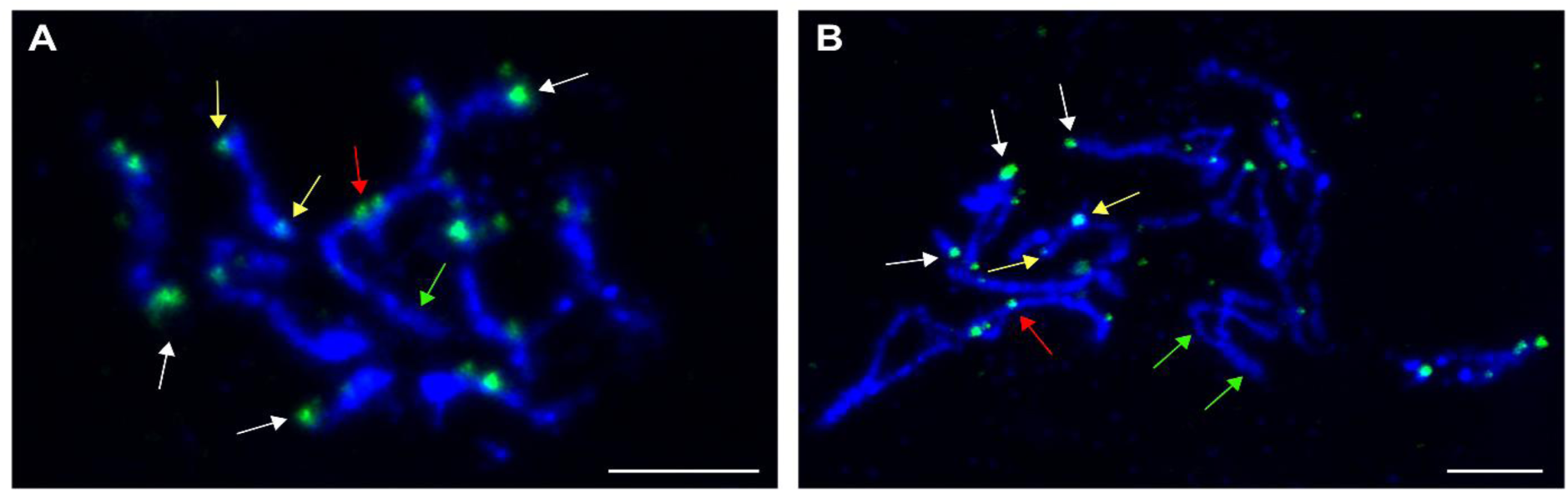
DNA FISH with the 16bp tandem repeat probe on *M. hapla* chromosomes in different chromosome condensation stages (A and B). Probe is labeled with FITC (green) and chromosomes are counterstained with DAPI (blue). Arrows point to hybridization signals localized at one (white arrows) or both (yellow arrows) chromosome ends. Red arrow indicates a chromosome where tandem repeats appear to be located internally. Green arrows point to chromosomes without hybridization signals. Size bar=5µm

Within the nematode order Rhabditida, the availability of improved chromosome-length assemblies led to the identification of seven ancient linkage blocks, known as Nigon elements (Gonzalez de la Rosa et al., 2021). These elements corresponded to six autosomes (A, B, C, D, E, N) and a sex chromosome (Nigon X), based on the co-localization of orthologous gene groups. These Nigon elements have been identified in multiple nematode species, including *C. elegans*, *Pristionchus pacificus*, and *Auanema rhodensis* (Tandonnet et al., 2019; Prabh & Rödelsperger, 2022; Rödelsperger, 2024). Using the previous *M. hapla* assembly, no clear Nigon assignments were identified, likely due to the highly fragmented nature of the assembly (Gonzalez de la Rosa et al., 2021). However, analysis of the current *M. hapla* assembly identified fragments of Nigon elements, with detectable, although highly fragmented, chromosomal segments. Nigon X, the ancestral sex chromosome, was not clearly identifiable (**Fig. 3).**

**Fig. 3:**
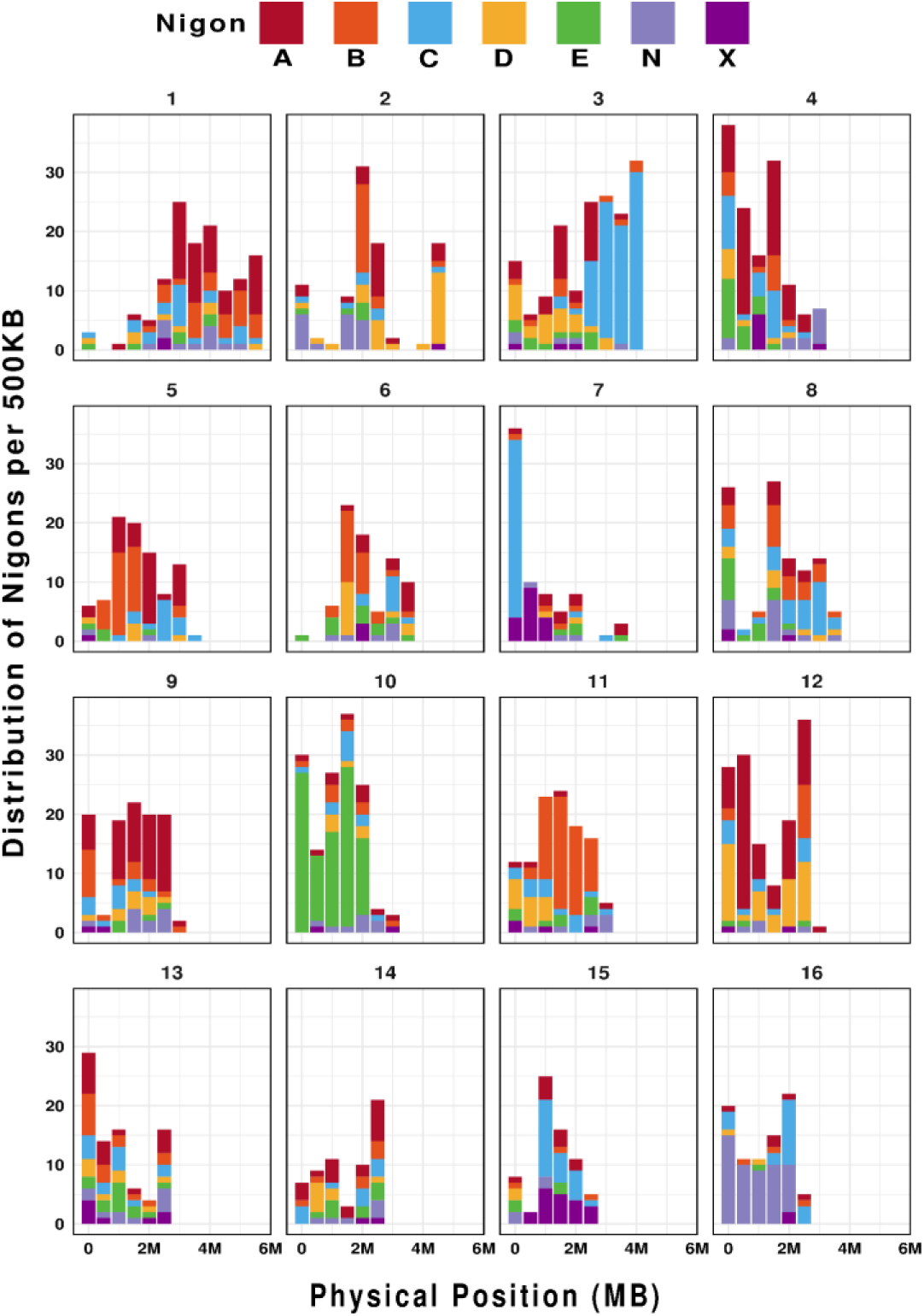
Distribution of Nigon elements along the length of *M. hapla* scaffolds. The X-axis represents the physical length of each scaffold, and the Y-axis represents the Nigon-defining loci per 500 Kb non-overlapping windows. The legend shows the color key for each Nigon Element from A through X.

### Genetic linkage map supports physical assembly and implies structural variation between nematode isolates

To assess whether our chromosome-length scaffolds correspond to true chromosomes, we produced a scaffold-based linkage map utilizing expressed sequence SNPs (eSNPs) from F2 lines derived from a cross between *M. hapla* strains VW9 and LM (Guo et al., 2017). Using the new assembly, we identified 789 SNP alleles flanking crossover events in each of 84 F2 lines (**Fig. 4**). The genome-wide recombination rate was high – 14.7 cM/Mb – compared to the 2.9 cM/Mb found in *C. elegans* (Rockman & Kruglyak, 2009). The recombination frequency profiles (Marey maps) for twelve scaffolds (S2, S3, S5, S6, S8, S9, S10, S11, S13, S14, S15, S16) are characterized by a high recombination rate in the chromosome arms and low recombination in the chromosome centers or tips (**Fig. 5**). This recombination pattern is similar to those described for *C. elegans*, *C. briggsae*, and *P. pacificus* (Rockman & Kruglyak, 2009; Ross et al., 2011). However, the remaining chromosomes showed intriguing deviations from the typical recombination profile. Most notably, S1 showed high recombination through the central region in addition to chromosomal arms, while S4 showed no recombination and S7, S12 and S13 showed low recombination. Intriguingly, segregation correlation in F2 lines suggested that S1 consisted of two linkage groups and that S4 and S12 alleles were on the same linkage group (**Supp. Fig. 5**). In addition, S13 showed highly skewed distribution of alleles in F2 lines favoring LM **(Fig. 4).**

**Fig. 4:**
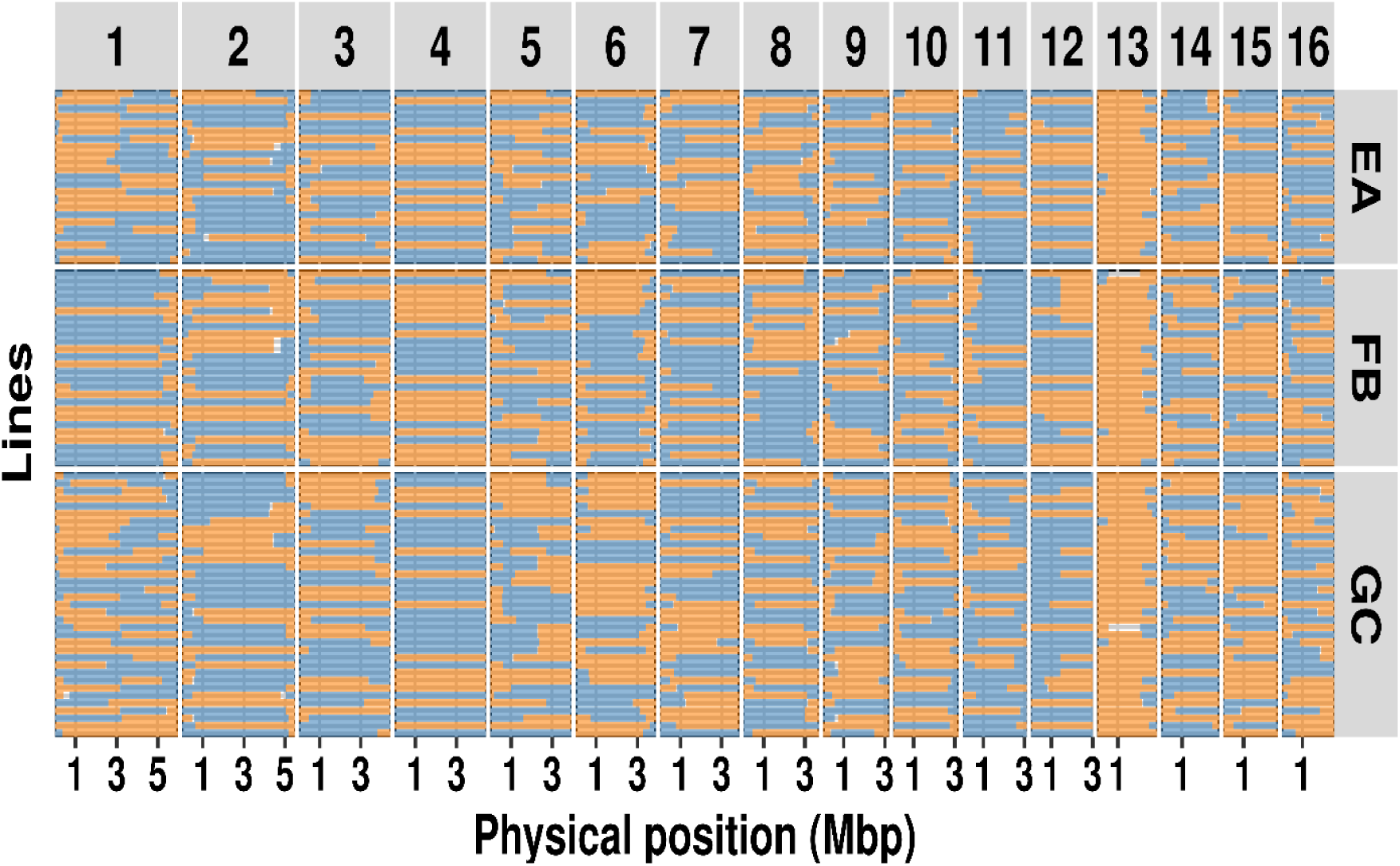
Recombination profile based on chromosome scaffolds. Allele present in each F2 line on scaffolds 1-16. Colors represent regions with VW9 (blue) and LM (orange) alleles, whereas white areas indicate insufficient data. Panels EA, FB, and GC on the right indicate the female from which the F2 lines were derived.

**Fig. 5:**
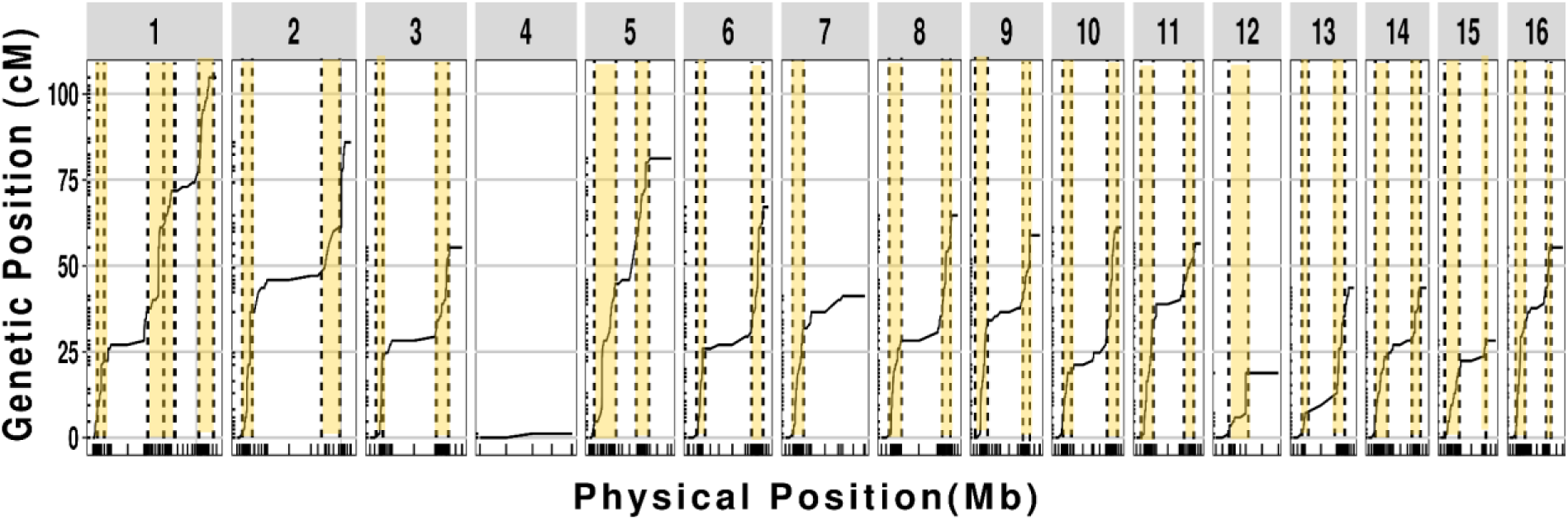
Recombination profile for each scaffold. Tick marks on the x axis indicate physical position on the scaffold and those on the y axis are the corresponding genetic positions based on SNP segregation in F2 lines. High recombination zones (HRZs) are highlighted in yellow. The number of scaffolds is depicted on top of each box.

We used change point analysis to further define the distribution and boundaries between regions with high and low recombination. This revealed sharp differentiation along the scaffolds between high recombination zones (HRZs), where rates ranged from 11 to 139 cM per Mb, and low recombination zones (LRZs) where rates were <5 cM/Mb **(Fig. 5; Supp. Table 2)**.

To test whether scaffold mis assembly was contributing to the anomalies that we observed in the scaffold-based linkage map, we produced a *de novo* classical genetic linkage map using segregation data from 458 SNPs in 93 RIL-like F2 lines from the previously described cross between *M. hapla* strains VW9 and LM (Guo et al., 2017). The resulting map contained 19 linkage groups (LGs). We then located the mapped SNPs on scaffolds in the new genome assembly to compare chromosome scaffolds to the genetic linkage groups (**Supp. Table 3**). For thirteen linkage groups, all SNPs were assigned to a single scaffold, supporting that these scaffolds represented full length chromosomes (**Table 1**). However, scaffolds S1, S2 and S13 were each divided into two genetic linkage groups. The most striking discrepancy was S1, which formed two linkage groups in the classical map.

**Table 1.** Alignment of *Meloidogyne hapla* VW9 scaffolds with genetic linkage groups. ^a^LG names are based on those assigned in previous work (Guo et al, 2017 ^a^; Thomas and Williamson, 2012^b^).

| Scaffold | Genetic Linkage Group |  |
| --- | --- | --- |
|  | VW9 X LM<sup>a</sup> | VW8 X VW9<sup>b</sup> |
| S1 | LG1a, LG1c | LG1 |
| S2 | LG2a, LG2c.15 | LG2, LG15 |
| S3 | LG4 | LG4 |
| S4 | LG14 | LG14, LG17 |
| S5 | LG10.12 | LG10, LG12 |
| S6 | LG2b | LG2 |
| S7 | LG6 | LG6 |
| S8 | LG3 | LG3 |
| S9 | LG1b | LG1 |
| S10 | LG16 | LG16 |
| S11 | LG5 | LG5 |
| S12 | LG13.17 | LG13 |
| S13 | LG9a, LG9b | LG9 |
| S14 | LG7 | LG7 |
| S15 | LG8 | LG8 |
| S16 | LG11 | LG11 |

As noted above, S1 showed anomalously high recombination in the center as well as on arms. Interestingly, the break between the linkage groups in the VW9xLM cross and the high central recombination occurred in the region of the internal palindrome corresponding to the telomeric repeat (**Fig. 1B**). To determine whether this joining was a scaffolding error, we examined nanopore reads and identified 2902 raw ultralong nanopore reads greater than 60 kb spanning the palindromic region of S1 (**Supp. Data 3**). Hence, the existence of a single molecule spanning this region is strongly supported. Additionally, we noted a difference between the recombination patterns in progeny from female FB, one of the three F1 females from which the RIL-like F2 lines derive showed no recombinants in the central region of S1 chromosome whereas progeny from the other two females (EA and GC) showed high apparent segregation in this region.

To further investigate the anomalies between scaffolds and linkage groups, we utilized a published genetic linkage map generated with segregation data of DNA polymorphisms in 183 F2 lines from a cross between *M. hapla* strains VW8 and VW9 (Thomas et al., 2012). We then located the mapped SNPs onto scaffolds in our new genome assembly. Ten linkage groups corresponded to single scaffolds (**Table 1**). However, even though VW9 was a parent in both crosses (VW9xLM and VW8xVW9), there were differences in the alignment of chromosome scaffolds to linkage groups for the two crosses. Notably, LG1 spanned all SNPs of S1 and S9, predicting it to be a very long linkage group. Also, LG2 contained all markers for scaffolds S2 and S6. Interestingly, the highly skewed marker segregation in favor of LM alleles seen for S13 in the VW9xLM cross was not observed in the VW8xVW9 cross. In addition, in the VW8x9 linkage map, S4 and S12 did not show repressed recombination. Also, the highly skewed segregation of S13 markers observed for the VW9xLM cross was not apparent in the VW8xVW9 cross. Since both genetic crosses used VW9 as one parent but a different second parent, the disparity between the chromosome scaffolds and linkage maps is likely due to structural differences between genomes of the three strains. For example, inversions, translocations or other genomic rearrangements (chromosome fusions, breakage) could result in the distorted segregation patterns that we observed. Other intriguing discrepancies between scaffolds and LGs remain to be explained but would likely require producing chromosome-length sequences of multiple *M. hapla* strains.

Together these results suggest that the differences between linkage maps and the new chromosomal *M. hapla* assembly are due to differences between the genome structure of VW9 and other parental strains. Therefore, the current assembly likely corresponds to a largely correct representation of the chromosomal DNA molecules in strain VW9.

### Characterization of predicted secreted protein gene repertoire of *M. hapla*

To annotate the newly generated VW9 genome, we utilized Iso-Seq data from mixed developmental stages, including eggs, J2, and females. This yielded 4,117,943 reads, which were clustered into 240,273 high-quality isoforms with an N50 of 2,018 bp. Additionally, RNA-seq data were obtained from nematode-infected roots of *Solanum lycopersicum cv* Moneymaker and *S. pimpinellifolium* cv. G1.1554 at five post-inoculation time points (5, 7, 10, 12, and 14 days after inoculation). The integration of these two datasets using BRAKER3 produced a high-quality structural annotation comprising 11,229 protein-coding genes. This gene count is lower than the 14,700 genes reported in the 2008 assembly (Opperman et al., 2008), which was likely due to the reliance in the previous study on abinitio predictions, which can overestimate gene numbers. The new annotation as assessed with BUSCO using Eukaryota_odb10 (Manni et al., 2021) showed 88.2% completeness for coding sequences with only 2.4% fragmentation **(Supp. Table 4).** Similarly, the annotation assessed using BUSCO using Nematoda_odb10 database showed 63.8% completeness for coding sequences with only 2.7% fragmentation. This represents a notable improvement over the 2008 assembly, which showed 78.5% completeness and 10.2% fragmentation with eukaryota_odb10, and 50.5% completeness with 4.1% fragmentation with nematoda_odb10 databases.

To identify genes likely involved in virulence (candidate effectors), we screened the 11,229 predicted proteins for the presence of secretion signal sequences (SignalP6.0; (Teufel et al., 2022) and the absence of transmembrane domains (DeepTMHMM; (Hallgren et al., 2022). This resulted in the identification of 1,258 genes encoding predicted secreted proteins (PSPs; **Supp. Data 4**). We then used the full set of 1,258 genes to perform ortholog searches across 71 nematode species using OrthoFinder. Of the 71 nematode species, 33 were plant parasitic nematodes and the rest were either free living or human parasitic nematodes. This analysis revealed that 1,172 of these PSPs had orthologs in at least one other species while 86 were unique to *M. hapla;* 675 *M. hapla* PSPs were conserved across all 71 nematode species; 401 were present only in RKNs; 3 conserved only with other PPNs; and 1 was shared with only one other nematode species (**Fig. 6A&B; Supp. Table 7**). Among the *M. hapla*specific PSPs, 56 were single-copy genes (singletons) and 30 present in multiple copies.

**Fig. 6.**
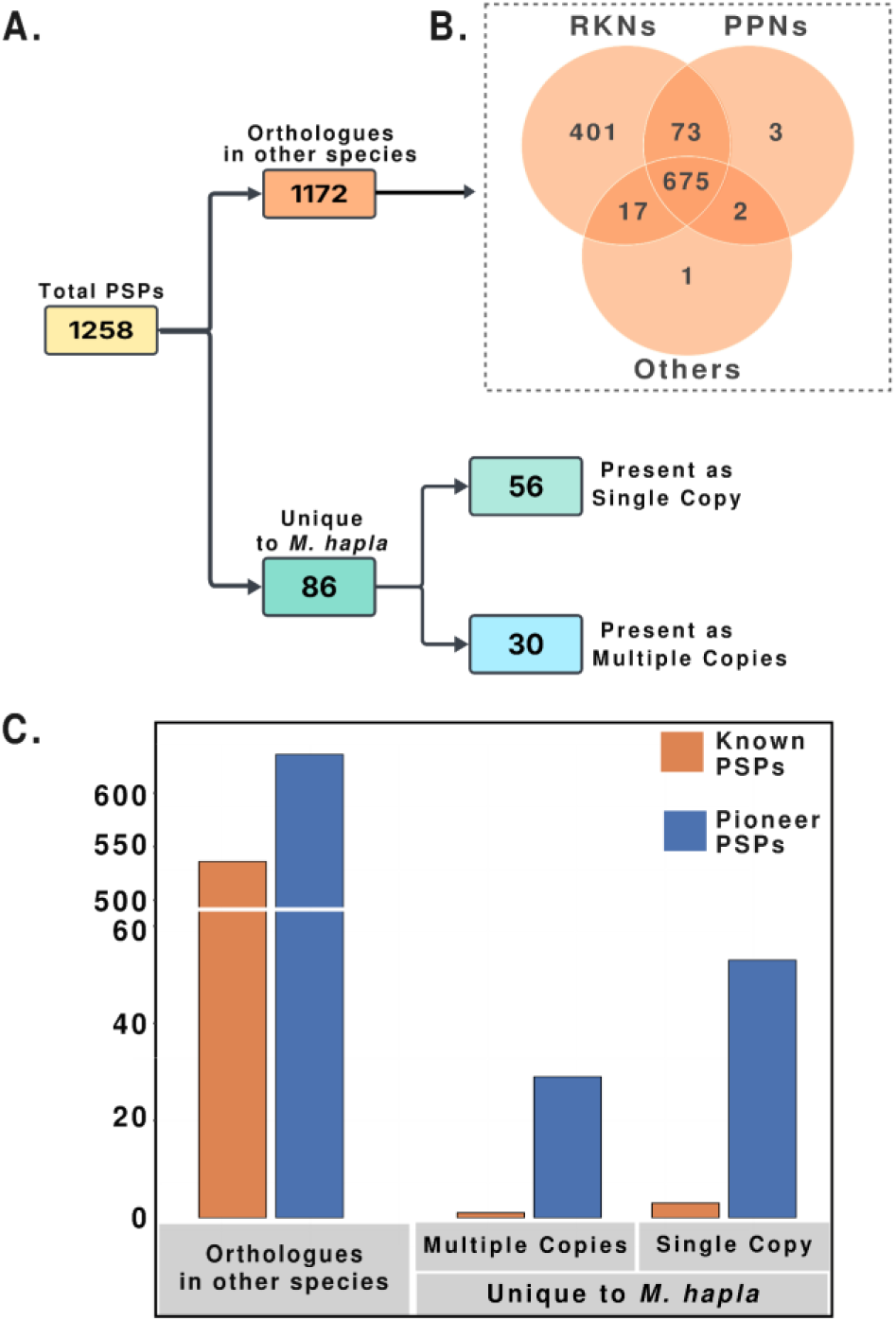
Classification of *M. hapla* PSPs. **A.** Total PSPs were characterized by the presence or absence of orthologs in a set of 71 nematode species. The identification of orthologs was done using Orthofinder. **B.** Venn diagram shows the PSPs that are shared across RKNs, PPNs and other nematode species. **C.** The distribution of PSPs into Known and Pioneer PSPs is shown for PSPs with orthologs in other nematode species and for those unique to M. hapla. (i.e., those unique to *M. hapla* having single or multiple copies.

Of the 1258 genes encoding PSPs in *M. hapla*, 540 contained known functional domains in Interpro and EggNOG databases, while the remaining 718 did not and are referred to as “pioneer PSPs” (**Supp. Table 5 & 6**). For the 1,172 PSPs that had orthologs in other nematodes, 536 contained known domains and 636 were pioneers (**Fig. 6C; Supp. Table 7**). Most of the PSP genes present only in *M. hapla* were pioneers. Of the single-copy PSPs unique to *M. hapla*, 53 had no known functional domains. Three encoded proteins with identifiable features, including a glycine-rich domain, a collagen triple helix repeat, and an SH3 domain. Similarly, among the 30 PSP genes unique to *M. hapla*, but present as multiple copies, only one had a known domain (a protein kinase).

Within the 540 PSP families containing a known functional domain, we identified multiple gene families previously shown to play a role in parasitism in other organisms (**Supp. Table 8**). These include several families of CAZymes–31 Glycoside Hydrolases (GH), 17 Glycosyl Transferases (GT), 7 Carbohydrate Esterases (CE), and 12 Pectate Lyases (PL) (Haegeman et al., 2011) (**Supp. Fig. 6, Supp. Table 9**). We further verified the annotation of these CAZymes through the dbCAN database (Yin et al., 2012) (**Supp. Table 10**). Previous studies suggest that horizontal gene transfers contributed to the acquisition of CAZymes in plant-parasitic nematodes (Haegeman et al., 2011). Using HGT analysis, we confirmed that a total of 20 CAZymes that include 7 GHs, 1 CE and all PLs have evidence of horizontal gene transfer (**Supp. Table 10**). Similarly, other PSPs (Alginate Lyases, Lytic transglycosylases, Fungal chitosanase, Protein Kinase domain containing protein, and Trypsin) also showed evidence of horizontal transfer (**Supp. Table 10**). Furthermore, consistent with previous phylogenetic studies that suggest pectate lyases expansion through gene duplication during *Meloidogyne* evolution (Opperman et al., 2008; Thomas et al., 2012), our analyses revealed patterns of clade separation and subsequent expansion across all four CAZymes families (**Supp. Fig. 7-9, Supp. Data 5**).

In our structural annotation pipeline, default filtering criteria were employed, which typically exclude very short open reading frames. In addition, the RNA-seq and Iso-seq datasets used in the annotation process were not optimal for identifying short transcripts, as the sequencing read lengths were often insufficient to detect and reconstruct these smaller genes with confidence. As a result, proteins shorter than 66 amino acids were likely not annotated, potentially omitting small, secreted proteins that may function as plant peptide mimics. To more comprehensively identify such candidates, we reanalyzed the 2008 genome annotation and identified 19 small PSPs. Among these, five had functional annotations, including a C2H2 zinc finger domain protein, a fructosyltransferase, a phosphotransferase system component, a bifunctional nuclease-like protein, and a tyrosinase copper-binding protein. The remaining 14 proteins lacked known functional annotations (**Supp. Table 11**). When mapped onto the updated *M. hapla* genome assembly, these PSPs were found to be distributed across multiple genomic regions (**Supp. Table 11**).

Next, we conducted a BLAST search for known plant peptide mimics, including C-terminally Encoded Peptides (CEPs), Inflorescence Deficient in Abscission (IDA) and Rapid Alkalinization Factors (RALF)—all previously described to modulate parasitism related responses in plants (Kim et al., 2018; Mishra et al., 2023; Zhang et al., 2020). We found that all 12 CEPs were located on S13, while IDA1 and IDA2 were located on S7 and S9, respectively. The three RALFs were located on S1, S5 and S10 (**Supp. Table 12**). No PSY peptide genes were detected, consistent with prior findings that these genes are restricted to Clade I *Meloidogyne* species (Yimer et al., 2023).

### Genes encoding Predicted Secreted Proteins (PSPs) are enriched in high recombination zones

To examine the genomic distribution of protein-coding genes, we compared their empirical cumulative distribution to a theoretical uniform distribution (**Supp. Fig. 10)**. Overall, genes appear evenly distributed, except in certain regions where anomalies coincide with unusual recombination patterns (**Fig. 5**). For instance, S1 separates into two genetic linkage groups in the VW9×LM cross, while S4 and S12 appear to be physically linked. In contrast, S7 contains only a single region with a markedly higher recombination rate (**Fig. 5**). This pattern of uniform gene distribution is consistent with observations in other holocentric species, such as spider mite (*Tetranychus urticae*) and *C. elegans*, where genes are generally evenly distributed along the chromosomes (Grbić et al., 2011; Spieth et al., 2018).

Given the rapid evolution of effectors under selection pressure in various plant pathogens (Fouché et al., 2018; Jagdale et al., 2021), we assessed whether PSP genes cluster in HRZs or LRZs (**Supp. Table 2**). A hypergeometric test for enrichment found that PSP genes were significantly enriched in HRZs (*p=2.5e-08*; **Supp. Table 13**). A scaffold-specific analysis found PSP enrichment on S2, S3, S14, and S16 while other scaffolds did not show significant differences in PSP distribution between HRZs and LRZs (**Fig. 7, Supp. Table 14**). In addition, annotated PSP families–Cysteine-rich secretory protein, Papain cysteine proteases, Aspartyl proteases, C-type lectin, peptidases, CAZymes, Catalases, Peroxidases, Astacins, Lipases and SXP/RAL2 family– were predominantly localized within HRZs (**Fig. 8**). One notable case is HRZ of S3, which exhibits the highest recombination rate in the genome at 139 cM/Mb (**Fig. 9A, Supp. Table 2).** This HRZ spans 340 KB, and contains 60 genes, 23 (∼38%) of which encode PSPs (**Fig. 9B & 8C)**. Among these, 16 are novel PSPs, while the remaining include known PSPs such as lysozyme, carboxylesterases, glycoside hydrolases, Galectin and Calycin (Lipocalin) (**Fig. 9C & 9D**).

**Fig. 7:**
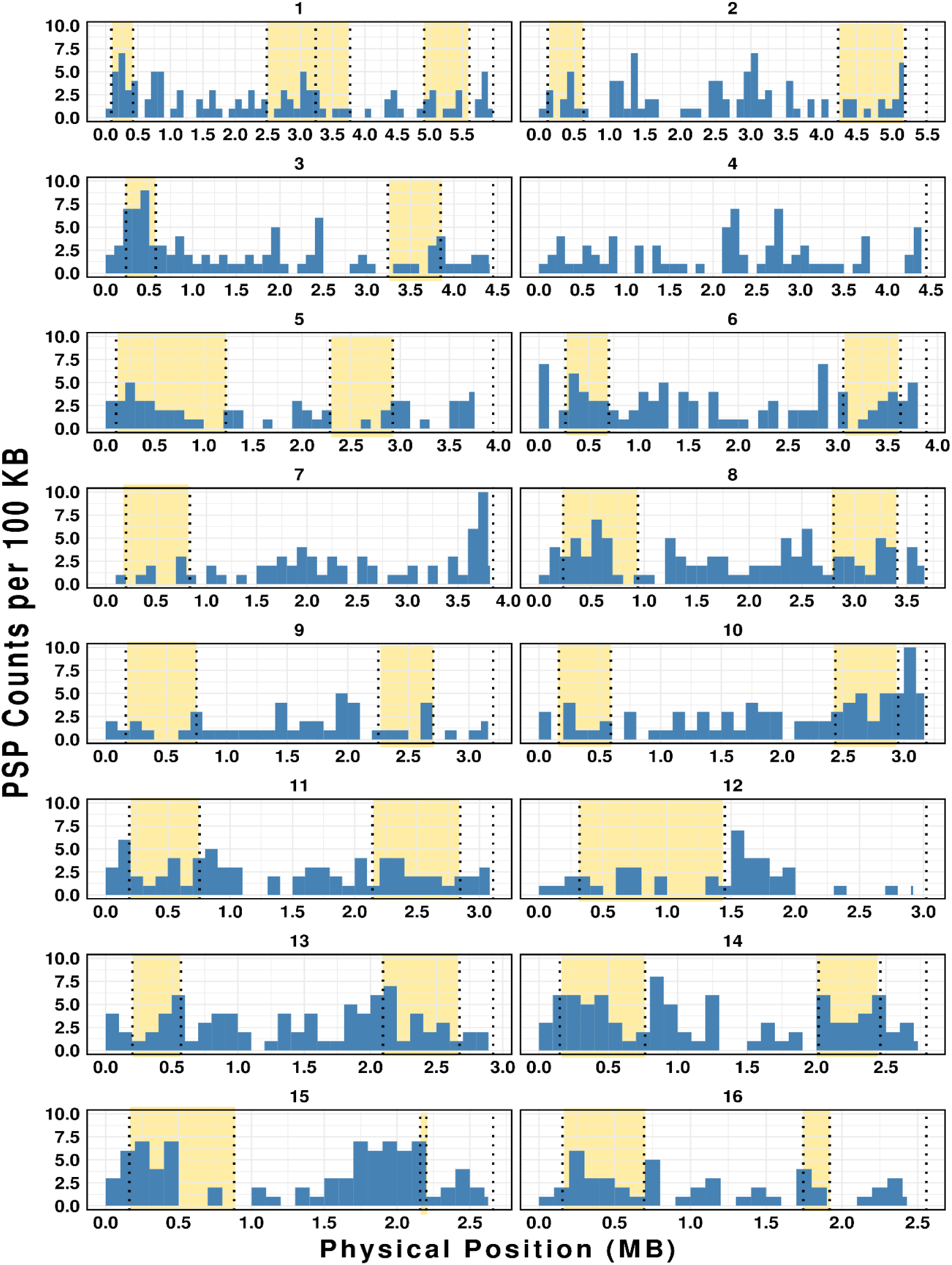
Distribution of PSP genes in the High Recombination Zones (HRZs). Each of the 16 scaffolds is divided into 100 kb bins ( X-axis). Y-axis represents the number of PSPs per bin. HRZ regions are highlighted in yellow. Genome-wide enrichment analysis shows significant enrichment of PSPs in the HRZs (Hypergeometric test, P-value=2.5e-08).

**Fig. 8:**
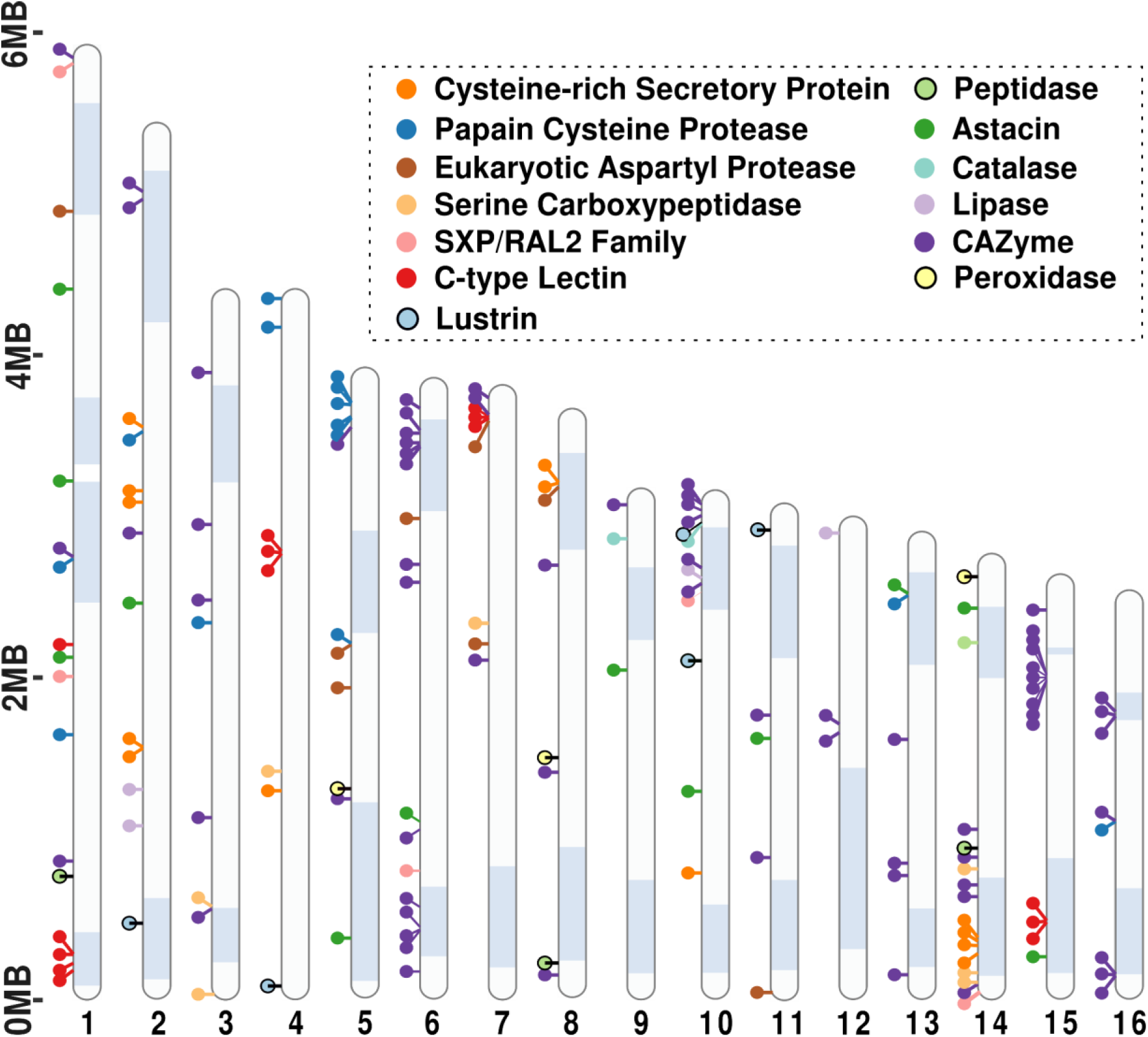
Distribution of known PSPs along the genome of *M. hapla*: Ideogram of well-known PSPs along the scaffolds of *M. hapla* genome. Each circle represents the PSPs, and they are pinned with respect to their positions on the scaffolds. The highlighted blue regions denote the HRZs.

**Fig. 9:**
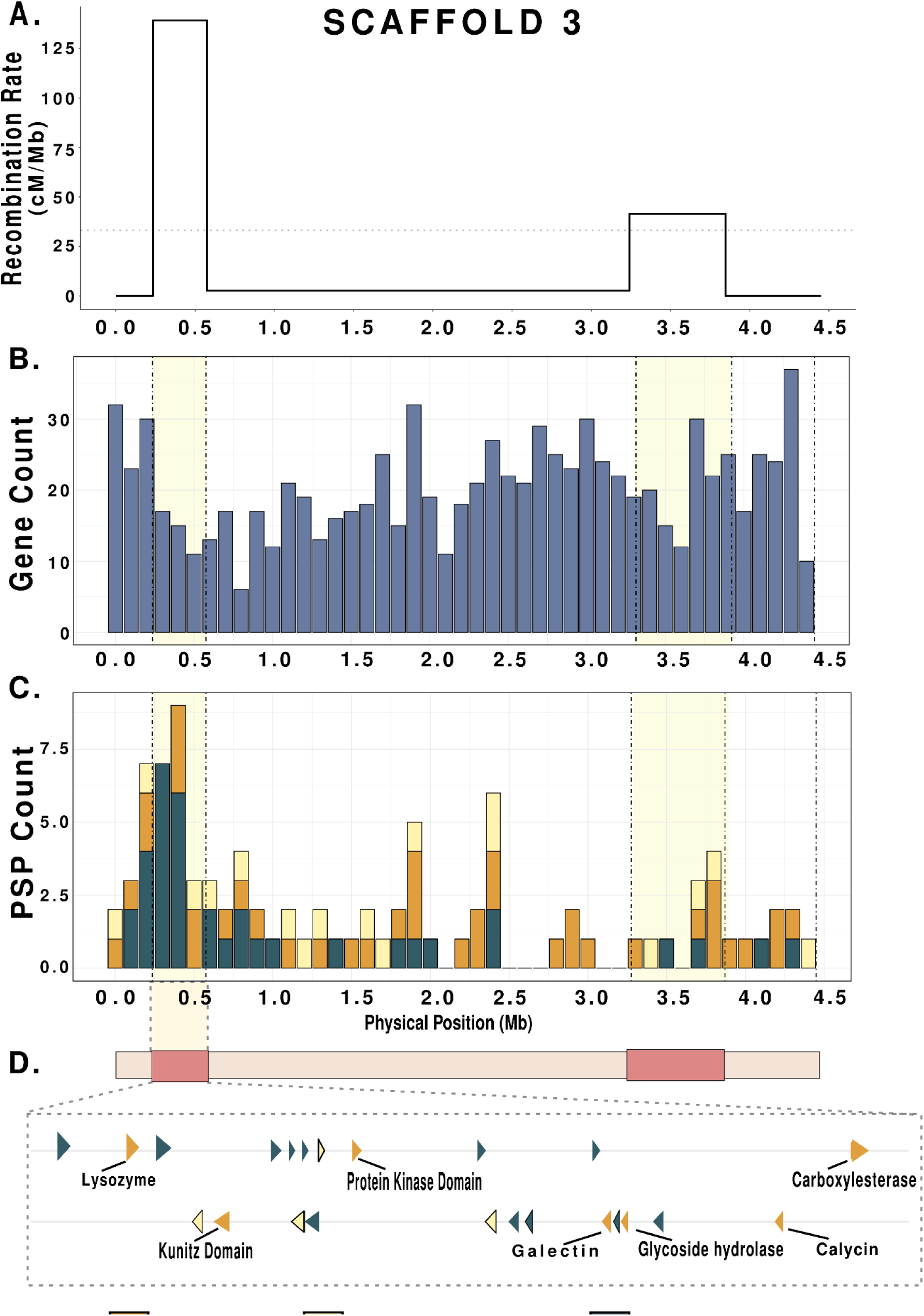
High recombination zones and gene distribution in Scaffold 3: **A.** Recombination rate (cM/Mb) of Scaffold 3 along its physical position (Mb) showing sharply defined HRZs. The dotted horizontal line indicates the average recombination rate for the whole chromosome. **B.** Histogram of Gene Count per 100 Kb along Scaffold 3. The highlighted regions denote HRZs. **C.** Stacked bar graph of PSP Count per 100 Kb along the physical positions of scaffold 3 where the highlighted regions denote HRZs. Each color represents functionally annotated PSPs, Novel PSPs shared with other Meloidogyne species and Novel PSPs that are unique to *M. hapla.* **D.** Highlights of the PSP genes found in HRZ of Scaffold 3. They are colored according to the ones shown in C. The arrows represent positive and negative strands. The known PSPs are labelled with their respective functional annotations.

In *C. elegans*, evolutionarily conserved genes are enriched in low recombination regions of chromosomes (Hillier et al., 2007). To assess whether *M. hapla* shows a similar pattern, we identified 5610 *C. elegans* orthologs from the Orthofinder analysis **(Supp. Data 6**). Of these, 4195 map to the LRZ and 1415 to the HRZ. An enrichment analysis revealed no overrepresentation of *C. elegans* orthologs in HRZs; however, these orthologs were significantly enriched LRZs.

## Discussion

Similar to other *Meloidogyne* species, *M. hapla* lacks telomerase and canonical telomere-associated proteins (Mota et al., 2023), suggesting that it relies on an alternative mechanism for chromosome-end maintenance. In this study, we present a chromosome-scale genome assembly for *M hapla* strain VW9, consisting of 16 scaffolds corresponding to the previously reported chromosome number for this strain. A notable finding of this new assembly is the identification of tandem repeats consisting of a consensus 16-mer sequence (CCCAAGGTTTAAAAGG) located at one or both ends of most scaffolds, and intriguingly, also in an interior region of Scaffold 1. These repeats show no recognizable homology to the more complex terminal repeats recently identified in Clade I *Meloidogyne* species (Dai et al., 2023; Mota et al., 2023) or to the terminal repeats in the diploid species *M. graminicola* (Dai et al., 2023). This lack of homology suggests that chromosome-end repeat motifs have significantly diversified among root-knot nematode species. Notably, most of the 16-mer repeats in our assembly occur at one scaffold end, although we cannot exclude the possibility that our assembly is incomplete or that additional relevant sequences remain undetected. The presence of repeats at only one end is also reminiscent of *C. elegans* pairing centers, which have been found to facilitate homologous pairing and crossover during meiosis (M. Li et al., 2024). Whether the terminal regions of *M. hapla* play a role in telomere maintenance, meiotic pairing, or both remain to be investigated.

Because *M. hapla* is capable of facultative sexual reproduction, it is possible to carry out controlled crosses and track allele segregation in RIL-like F2 lines. These data provide a valuable resource for studying chromosome structure and parasitism-related adaptation in plant-parasitic nematodes. The recombination patterns of individual F2 lines along the majority of scaffolds in *M. hapla* resemble those observed in *C. elegans*, with crossovers primarily localized to the arms. However, the overall recombination rate in *M. hapla* is roughly five times higher than in *C. elegans*, where meiotic crossovers are restricted to one per chromosome per meiosis (Hillers & Villeneuve, 2003; Yu et al., 2016; Rockman & Kruglyak, 2009). The *M. hapla* genetic map suggests that the same per-chromosome crossover limit may also apply to this species. Therefore, its higher recombination rate could be primarily due to its higher chromosomes number and smaller genome size. Recombination in *M. hapla* is localized to more narrowly defined regions with sharp boundaries, resulting in very high local recombination rates. Intriguingly, electron microscopic studies have described the presence of synaptonemal complexes and distinctive electron-dense “recombination nodules” in oocytes of the meiotic race of *M. hapla* (Goldstein & Triantaphyllou, 1978).

Comparison between genome assembly scaffolds for *M. hapla* strain VW9 and genetic maps derived from F2 crosses (VW9×LM and VW8×VW9) indicates genome rearrangement differences between the three nematode strains **(Table 2)**. The parental strains were derived from geographically distinct field isolates: VW9 from California, VW8 from the Netherlands, and LM from France (Liu & Williamson, 2006; Wang et al., 2010). Reduced recombination observed in S4, S7, and S12 in the VW9xLM cross **(Fig. 4**; **Fig. 5)** is typical of crosses between parents with genome differences, such as translocations and inversion. S1 exhibits particularly intriguing segregation patterns. Both our assembly and ultra-long sequencing reads support S1 as a single chromosome. However, F2 progeny from the VW9×LM cross display high recombination in the central region of this chromosome for progeny from two of three F1 females (females EA and GC), whereas progeny from the third female (female FB) exhibit no recombination (**Fig. 4**). This finding suggests that S1 may exist as a single chromosome in female FB, but as two separate chromosome-like entities in females EA and GC. Interestingly, the classical linkage map from the VW8×VW9 cross indicates that scaffold S1 is linked with another scaffold (S9), forming a single chromosome. Such variability aligns with earlier global surveys that reported differences in chromosome numbers and modes of reproduction among *M. hapla* field isolates (Triantaphyllou & Hirschmann, 1980). Indeed, a cross between females with 15 chromosomes and males with 17 chromosomes produced viable progeny possessing more than 15 chromosomes, suggesting chromosome number fluidity within this species (Triantaphyllou, 1966). Holocentric chromosomes, like those found in *M. hapla*, may be particularly tolerant to fragmentation and fusion events compared to monocentric chromosomes. Additionally, a non-canonical mechanism for chromosome-end maintenance might further facilitate such genomic plasticity.

Some, but not all, segregation anomalies that we found in F2 line segregation patterns are consistent with inversions, translocations, or other chromosome structure differences between the parental lines. In another intriguing observation, we found that for F2 lines of the VW9xLM cross, alleles on Scaffold 13, are strongly skewed towards LM (**Fig. 4),** but this bias is not observed in the VW8xVW9 F2 lines. This strongly biased segregation pattern matches those found with toxin-antidote elements that have been observed in multiple species, including the nematode species *C. elegans* and *P. pacificus* (Seidel et al., 2008; Burga et al., 2019; Zdraljevic et al., 2024). Occurrence of such elements hamper overall reproduction success and strongly favor the presence of the toxin-antidote allele in the offspring. In this case, the data is consistent with the presence of a toxin-antidote element in strain LM. Insight into other observed segregation anomalies will require further analysis and additional genetic crosses.

In our annotation analysis, we focused on PSPs, as these likely include most genes involved in parasitism. Our ortholog analysis revealed considerable overlap between the PSPs of *M. hapla* and other *Meloidogyne* species (**Fig. 6)**. Most PSPs with known protein motifs, including CAZymes and genes acquired via HGT, are shared across RKN species. This pattern suggests that many HGT events occurred before the recent speciation of these nematodes. Furthermore, over half of the PSP genes that *M. hapla* shares with other RKN spp have no identified functional motifs, and nearly all PSP genes that are unique to *M. hapla* are pioneers. We speculate that these unique pioneer PSPs may have originated via *de novo* gene birth or represent orphan genes derived from mechanisms such as gene duplication.

The availability of genetic segregation data provides a unique opportunity to investigate the relationship between recombination rates and the genomic distribution of parasitism genes in meiotic RKN species. In other pathogenic organisms, effector genes are often enriched in genomic regions associated with high recombination. For example, recombination hotspots in the fungal pathogen *Blumeria graminis* harbor effector genes (Müller et al., 2019). Similarly, in *Fusarium graminearum* and *Zymoseptoria tritici*, effectors are enriched in chromosome regions near sub-telomeres or on accessory chromosomes, regions characterized by higher recombination rates and described as having a “two-speed” genome architecture (Croll et al., 2015; Laurent et al., 2017). Our genome-wide analysis of PSP distribution found that these genes are generally enriched in HRZs (**Fig. 7**). However, a scaffold-specific analysis showed significant PSP enrichment in HRZs for only four out of 16 scaffolds indicating that the enrichment of PSPs is unevenly distributed across scaffolds (**Fig. 8**; **Fig. 9; Supp. Table 14)**. By contrast, conserved orthologs shared with *C. elegans* showed enrichment in LRZs. Thus, enrichment in HRZs appears to be specific to PSPs rather than a general feature of gene localization in *M. hapla*.

Double-strand DNA breaks produced during meiosis have been shown to promote genomic variation in humans (Hinch et al., 2023). In line with this, we hypothesize that the enrichment of PSPs in HRZs may facilitate diversification, enabling these genes to rapidly evolve, evade host recognition, and overcome host resistance. Interestingly, Clade I *Meloidogyne* species, which reproduce asexually and lack meiosis, also exhibit broad host ranges and can evolve to avoid host resistance. These asexual species are of hybrid origin and carry two divergent subgenomes (Blanc-Mathieu et al., 2017; Szitenberg et al., 2017; Dai et al., 2023), suggesting that their ability to acquire new specificity may be due to interactions between the subgenomes (Szitenberg et al., 2017). Thus, the diploid, meiotic *M. hapla* and the asexual Clade I species likely achieve adaptability through fundamentally different genomic mechanisms. Understanding how recombination and genome architecture drive PSP evolution can provide new insights into host-parasite co-evolution as well as support development of control strategies for nematode infestations. Future studies will focus on comparative genomics and functional analysis to identify evolutionary processes underlying PSP diversification in these important plant pathogens.

## Materials and Methods

### High molecular weight DNA extraction

*Meloidogyne hapla* strain VW9 (Liu & Williamson, 2006) was propagated on tomato (*Solanum lycopersicum*) cultivar VFNT cherry in a UC Davis Greenhouse facility. Nematode eggs (∼1 million) were collected from roots then cleaned by sucrose floatation method and flash frozen in liquid nitrogen. High molecular weight (HMW) gDNA extraction was carried out at UC Davis Genome Center. Two ml of lysis buffer containing 100 mM NaCl, 10 mM Tris-HCl pH 8.0, 25 mM EDTA, 0.5% (w/v) SDS and 100 µg/ml Proteinase K was added to the tube containing frozen eggs. The samples were mixed with gentle pipetting and homogenized at room temperature overnight. The lysate was then treated with 20 µg/ml RNAse at 37 °C for 30 minutes. The lysate was cleaned with equal volumes of phenol/chloroform using phase lock gels (Quantabio Cat # 2302830). The DNA was precipitated from cleaned lysate by adding 0.4X volume of 5M ammonium acetate and 3X volume of ice-cold ethanol. The DNA pellet was washed with 70% ethanol twice and resuspended in an elution buffer (10mM Tris, pH 8.0). The purity of gDNA was assessed using NanoDrop spectrophotometer and 260/280 ratio of 2.0 and 260/230 of 2.29 were observed. For PacBio HiFi, DNA yield was 8 µg as quantified by Qubit 2.0 Fluorometer (Thermo Fisher Scientific, MA). The integrity of the HMW gDNA was verified on a Femto pulse system (Agilent Technologies, CA) where 68% of the DNA was found in fragments above 100 Kb.

### PacBio HiFi sequencing for genome assembly

The HiFi SMRTbell library was constructed using the SMRTbell prep kit 3.0 (Pacific Biosciences, Menlo Park, CA, Cat. #102-182-700) according to the manufacturer’s instructions. HMW gDNA was sheared to a target DNA size distribution between 15-20 kb using Diagenode’s Megaruptor 3 system (Diagenode, Belgium; cat. B06010003). The sheared gDNA was concentrated using 1X of SMRTbell cleanup beads provided in the SMRTbell prep kit 3.0 for the repair and a-tailing incubation at 37°C for 30 minutes and 65°C for 5 minutes, followed by ligation of overhang adapters at 20°C for 30 minutes, clean-up using 1X SMRTbell cleanup beads, and nuclease treatment at 37°C for 15 minutes. The SMRTbell library was size selected using 3.1X of 35% v/v diluted AMPure PB beads (Pacific Biosciences, Menlo Park, CA; Cat. #100-265-900) to progressively remove SMRTbell templates <5kb. The 15 – 20 kb average HiFi SMRTbell library was sequenced at UC Davis DNA Technologies Core (Davis, CA) using one 8M SMRT cell (Pacific Biosciences, Menlo Park, CA; Cat #101-389-001), Sequel II sequencing chemistry 2.0, and 30-hour movies on a PacBio Sequel II sequencer. This way, we generated 232,679 sequences with an average length of 14,391 base pairs.

### Nanopore sequencing for genome assembly

For Nanopore sequencing, the sequencing libraries were prepared from 1.5µg of high molecular weight DNA using a ligation sequencing kit SQK-LSK114 (Oxford Nanopore Technologies, Oxford, UK). The manufacturer’s library preparation protocol was followed apart from extended incubation times for DNA damage repair, end repair, ligation and bead elution. For sequencing, the PromethION device P24 was used. Thirty fmol of the final library from the sample was first loaded on the PromethION flow cell R10.4.1 (Oxford Nanopore Technologies, Oxford, UK) and run was set up using PromethION MinKNOW 22.12.5 for 17 hours. Data was base called live during sequencing with super-high accuracy mode using ONT-guppy-for-promethion 6.4.6. This way, we generated 584,625 reads with an average length of 27,317 base pairs.

### RNA extraction for IsoSeq sequencing

Total RNA was extracted from three life stages of *M. hapla*: eggs, newly hatched juveniles, and female nematodes dissected from tomato roots 21 days post infection. RNAeasy mini kit was used as per the manufacturer’s instruction for RNA extraction (Qiagen, catalog nr. 74106). To eliminate genomic DNA from RNA samples, TURBO DNAase (Life Technologies, AM1907) treatment was administered. The concentration and purity of the extracted RNA were determined using Nanodrop One^C^ Microvolume UV-Vis Spectrophotometer, while RNA integrity and quality were evaluated using bioanalyzer.

### IsoSeq sequencing for genome annotation

cDNA and SMRTbell library were constructed using SMRTbell® prep kit 3.0 (PN 102-396-000 REV02 APR2022) according to manufacturer’s instructions. The cDNA was synthesized using NEBNext® Single Cell/Low Input cDNA Synthesis & Amplification Module (New England Biolabs, Ipswich, MA, Cat. #E6421L). After fifteen cycles of PCR for cDNA amplification, the cDNA was purified using 0.86X SMRTbell cleanup beads. Subsequently, the cDNA libraries derived from each sample were pooled, and used for SMRTbell library construction using the SMRTbell prep kit 3.0 (Pacific Biosciences, Menlo Park, CA, Cat. #102-182-700) with the following enzymatic steps: repair and a-tailing, ligation of overhang adapters, and nuclease treatment. The Iso-Seq SMRTbell library was sequenced at UC Davis DNA Technologies Core (Davis, CA) using one 8M SMRT cell (Pacific Biosciences, Menlo Park, CA; Cat #101-389-001), Sequel II sequencing chemistry 2.0, and 24-hour movies on a PacBio Sequel II sequencer.

### RNA extraction for short-read time-series RNAseq

Total RNA from the early stage of nematode infection was extracted from galls of tomato-infected roots at five different time points post inoculation. Fourteen days after sowing, tomato plants of *Solanum lycopersicum* cv. Money Maker and *Solanum pimpinellifolium* were inoculated in-vitro with either 0 or 200 surface-sterilized J2s of *M. hapla* strain VW9 (as described in (Verhoeven et al., 2023)). At 5, 7-, 10-, 12-, and 14-days post inoculation (dpi), all galls present in a single square dish-plate were dissected and flash frozen in liquid nitrogen for RNAseq. In non-infected plants, to reduce the variation in development, roots segments adjacent to the roots of infected plants were dissected as control. Per time points, two to three technical replicates were taken per plant per nematode treatments.

Liquid nitrogen flash-frozen root galls were smashed and homogenized using Tissuelyzer (Qiagen, Hilden). RNA extraction was performed using the Maxwell 16 LEV-plant RNA kit (Promega) following the manufacturer’s instruction. After isolation, RNA concentration and purification was evaluated using NanoPhotometer^®^ and spectrophotometer (IMPLEN, CA, United States). RNA integrity checking, library preparations, RNA sequencing and quality filtering was done using BGISEQ-500 at BGI TECH SOLUTIONS (Hongkong) with at least 50 million clean paired-end reads of 150bp per sample.

### HiC sequencing for scaffolding

Chromatin conformation capture data was generated using a Phase Genomics (Seattle, WA) Proximo Hi-C 4.5 kit, which is a commercially available version of Hi-C Protocol (Lieberman-Aiden et al., 2009). For library preparation, approximately 500 mg tissue was finely chopped and then crosslinked for 15 minutes at room temperature with end-over-end mixing in 1 mL of Proximo crosslinking solution. Crosslinking reaction was terminated with a quenching solution for 20 minutes at room temperature again with end-over-end mixing. Quenched tissue was rinsed once with 1X Chromatin Rinse Buffer (CRB). The tissue was transferred to a liquid nitrogen-cooled mortar and ground to a fine powder. Powder was resuspended in 700 μL Proximo Lysis Buffer 1 and incubated for 20 minutes with end-over-end mixing. A low-speed spin was used to clear the large debris and the chromatin containing supernatant transferred to a new tube. Following a second higher speed spin, the supernatant was removed and the pellet containing the nuclear fraction of the lysate was washed with 1X CRB. After removing 1X CRB wash, the pellet was resuspended in 100 μL Proximo Lysis Buffer 2 and incubated at 65°C for 15 minutes.

Chromatin was bound to Recovery Beads for 10 minutes at room temperature, placed on a magnetic stand, and washed with 200 μL of 1X CRB. Chromatin bound on beads was resuspended in 150 μL of Proximo fragmentation buffer and 2.5 μL of Proximo fragmentation enzyme added and incubated for 1 hour at 37°C. Reaction was cooled to 12°C and incubated with 2.5 μL of finishing enzyme for 30 minutes. Following the addition of 6 μL of Stop Solution, the beads were washed with 1X CRB and resuspended in 100 μL of Proximo Ligation Buffer supplemented with 5 μL of Proximity ligation enzyme. The reaction was incubated at room temperature for 4 hours with end-over-end mixing. To this volume, 5 μL of Reverse Crosslinks enzyme was added and the reaction incubated at 65°C for 1 hour. After reversing crosslinks, the free DNA was purified with Recovery Beads and Hi-C junctions were bound to streptavidin beads and washed to remove unbound DNA. Washed beads were used to prepare paired-end deep sequencing libraries using the Proximo Library preparation reagents. HiC sequencing was performed on Illumina (Novaseq) which generated over 52 million read pairs.

### Illumina DNAseq for polishing

DNA extraction from eggs was performed using the CTAB method as described in Dai et al., 2023. KAPA HyperPrep Kit was used according to the manufacturer’s protocol for library preparation. This involved processes such as enzymatic fragmentation, end-repair, A-tailing, and Illumina-compatible adapter ligation followed by PCR amplification and cleanup using magnetic beads. The final libraries were quantified using Qubit and assessed for size distribution using an Agilent Bioanalyzer. Consequently, the libraries were sequenced on the Illumina NovaSeq X platform using paired-end 150 bp reads. The library preparation and sequencing were performed by Novogene, USA. This way we generated 27 million read pairs.

### Genome profiling of PacBio HiFi reads

The genome profiling of the raw reads generated with PacBio HiFi was conducted using Jellyfish v.2.3.0, Genomescope2 and Smudgeplot0.2.5 (Marçais & Kingsford, 2011; Ranallo-Benavidez et al., 2020). K-mer frequencies for the raw reads were determined using Jellyfish with parameters set at “-m 21 -s 1000000000”. To deduce the lower and upper coverage range for these reads, “smudgeplot.py cutoff” was utilized. Subsequently, Smudgeplot was generated by using these lower and upper coverage values. Genomescope analysis was performed using default settings with the histogram data generated by Jellyfish as an input.

### Generation of draft assemblies

Multiple draft assemblies of the *M. hapla* genome were made using a combination of sequencing reads using various assembly software. For the final draft assembly that was used to make the final chromosome-scale assembly, we used Hifiasm v.2.0.0 (Cheng et al., 2021). First, we used Canu to correct the ONT reads to ensure the higher accuracy of ONT reads (Koren et al., 2017). After this, we incorporated PacBio HiFi, Canu corrected ONT reads, and raw Hi-C reads into Hifiasm to make a haplotype resolved assembly. The primary assembly generated using this method was polished using Illumina sequencing data in five cycles with Pilon v.1.23 using a Snakemake pipeline (See GitHub) (Walker et al., 2014). This corrected 222 bases, introduced 154 insertions, and removed 255 bases. Further, we used Meryl v.1.0 and Merqury v.1.3 to evaluate the quality of the genome assembly and Blobtools2 to check for any contamination in the genome (Miller et al., 2008; Challis et al., 2020; Rhie et al., 2020). For Blobtools, we followed the protocol outlined in this <u>Github repository</u> in which, we used reference proteomes (release:2024_04) from the UNIPROT database for diamond BLASTx, and Blast nt database (2.14), in addition to minimap2 v.2.26-r115 and samtools v.1.17 (Buchfink et al., 2021; Griffin, 2023/2023; H. Li, 2013, 2018). For BUSCO analysis, we used eukaryota_odb10 and nematoda_odb10 to check the quality of genome, CDS and protein sequences.

### Chromosome-scale assembly

We used HiC sequencing of VW9 strain to scaffold the draft genome into a chromosome scale assembly of the *M. hapla* genome. HiC data were generated by Phase Genomics using their Proximo HiC Kit Protocol with four cutter restriction enzymes: DpnII, DDel, Hinfl and Msel. Accordingly, we obtained the restriction sites and ligation junctions of these enzymes and generated restriction enzyme cut sites using the *generate_site_positions.py* script within the Juicer v.1.6. The information of the restriction sites was input within the Juicer v.1.6, and it was run with the early exit parameter (Durand et al., 2016). The HiC reads were aligned with the draft genome assembly with bwa alignment v.0.7.17-r1188 (H. Li, 2013). This was followed by quality control and filtering where low quality and chimeric reads were filtered out. The filtered aligned reads were merged into merged_sort.bam file. After the removal of duplicates from the merged file, merged_nodups.txt file was obtained which consisted of interaction matrices. The information obtained from Juicer was used to produce scaffolded assembly using 3D-DNA v.180114 (Dudchenko et al., 2017). 3D-DNA uses the interaction matrices to group the contigs into scaffolds. The initial scaffolded output was used to produce a draft contact map. This contact map was manually curated in Juicebox v.1.11.08 (Durand et al., 2016). The manual curation involved minor correction of contig positions according to the contact map. The manually curated assembly was then reviewed and finalized using run-asm-pipeline-post-review.sh script from 3D-DNA pipeline and final HiC contact map was made with the sizes of scaffolds in descending order.

### Nigon elements

The output of BUSCO run ‘full_table.tsv’ was used as an input to visualize the Nigon elements using vis_ALG ( https://github.com/pgonzale60/vis_ALG) (Gonzalez de la Rosa et al., 2021).

### Structural and functional annotation

Before structurally annotating the genome, the draft chromosome scale assembly was masked using RepeatModeler v.2.0.5 and RepeatMasker v4.1.5 (Flynn et al., 2020; Smit et al., 2021). For Iso-Seq pre-processing, we used isoseq3 v.4.0.0 which used ppbam v.2.4.99, pbcopper v.2.3.99, pbmm2 v.1.11.9, minimap2 v.2.15, parasail v.2.1.3, boost v.1.77, htslib v.1.17 and zlib v.1.2.13 (Ren et al., 2023). First, we clustered the demultiplexed Full-Length-Non-Chimeric (FLNC) reads using isoseq cluster tool. The high quality clustered and polished reads were then mapped to the genome using pbmm2 align with preset ISOSEQ parameter. After this, the reads were collapsed using isoseq3 collapse with parameters *–do-not-collapse-extra-5exons, –max-5p-diff5’ and –max-3p-diff5’*. These parameters prevent collapsing of isoforms that have extra 5’ exons and allow up to 5 base pairs difference at both 5’ and 3’ ends of the sequences ensuring that transcript diversity was preserved.

For structural genome annotation, we used BRAKER3 installed from its long-read branch (Gabriel et al., 2024). In the first run, BRAKER3 incorporated short read RNA-seq reads to guide the annotation. In the second run, BRAKER3’s long read protocol was applied during which GeneMarkS-T predicted the protein-coding regions from the long-read transcripts. Finally, TSEBRA was used to combine the data from both short-read and long-read evidence to produce a finalized structural annotation. For functional annotation of the predicted protein coding genes, we used BLAST2GO from OmicsBox v.3.3.2, InterproScan v.5.72-103.0 and Eggnog v.2.1.12 with eggnog database v.5.02 and novel family database v.1.0.1 (P. Jones et al., 2014; Cantalapiedra et al., 2021).

### Genetic map based on eSNPs of *M. hapla*

We produced a scaffold-based recombination profile using expressed sequence SNPs (eSNPs) previously identified between 98 RIL-like F2 lines of *M. hapla* produced by crossing strains VW9 and LM (Guo et al., 2017). The transcriptome data in that study was produced using RNA extracted from *Medicago truncatula* root galls infected with each RIL-like line and was made available from GEO under accession numbers PRJNA229407 and SRP078507. We used hisat2 (v. 2.2.1) to align the reads to the newly assembled chromosome-level genome. Thereafter reads were sorted, duplicate reads were removed, and the reads were indexed using Samtools (v. 1.14). Quality and alignment statistics were generated using fastqc (v. 0.12.1), Samtools, and mosdepth (v. 0.3.3). We used multiqc (v. 1.14) to inspect the qualities per sample. Next, variants were called using bcftools (v. 1.16) with standard settings except (ploidy = 2, keep_alts = true, min_mq = 20, min_bq = 13, min_idp = 5, max_idp = 500, min_qual = 20, min_ad = 2). This generated a file with 58,114 variants, which were further quality filtered using custom scripts in R (v. 4.2.2).

Next, we filtered the variants based on occurrence over the genotypes. We expressed variant occurrence as alternative allele frequencies (fraction of alt calls over total calls). When filtering for at least one clear alternative allele call we found 15,248 sites with at least one alt call. We then removed sites with more than 30% of samples uncalled, leaving 8,455 variants that were evenly distributed over the chromosomes. We called crossovers based on changepoint analysis using the changepoint package (v. 2.2.4) in R. Per RIL per chromosome, we first interpolated the uncalled sites based on the median genotype of the adjacent five variants before and after the uncalled variant. Subsequently, we used cpt.mean with the method BinSegto identify crossovers. The crossovers were visually inspected for accuracy.

Guo et al (2017) reported that some of the RIL-like lines appeared to be heterozygous. We confirmed this and therefore removed data from EA18, EA30, FB15, FB22, GC4, GC5, GC6, GC31, and GC45 before further analysis. Furthermore, for five lines the coverage by RNAseq was very low, requiring too much imputation to be of use (EA16, FB19, 12_GC14, GC36, and GC46). This left us with genetic data for 84 RIL-like lines. The set of markers for these lines was pruned for informative markers (markers at either side of a recombination event). The genotype at the ends of the chromosomes were interpolated from the first and last called marker per RIL. In total, this yielded a set of 789 markers for the 84 RILs (**Supp. Table 15**).

For recombination analysis we first calculated the genetic distances in centimorgans using a custom R script. We analysed the physical versus the genetic distance using changepoint package in R, we then identified transition points between the domains for each linkage group by performing a change-point analysis using the binary segmentation method (Killick & Eckley, 2014).

### Integration of genetic linkage maps with chromosome scale assemblies

To produce a framework genetic linkage map for chromosome scale assembly, we used segregation data from 458 SNPs in 93 RIL-like F2 lines produced from previously described cross between *M. hapla* strains VW9 and LM (Guo et al., 2017). Previously, this segregation data was used to produce a *de-novo* genetic map using the previous genome assembly of *M. hapla* obtaining 19 linkage groups (Opperman et al., 2008; Thomas et al., 2012). To identify the SNP positions in our genome assembly, we extracted sequences spanning 250 bp both upstream and downstream of each of the 458 SNPs using Bedtools v.2.31 (Quinlan & Hall, 2010). These sequences were subjected to a BLAST against chromosome-scale assembly. We then manually located the positions of the SNPs in the chromosome scale assembly which was used to align chromosome length scaffolds to the genetic linkage groups. The same strategy was used to anchor 182 genetic markers developed from an earlier genetic cross in which *M. hapla* strain VW8 as female parent was crossed with VW9 to the current genome assembly (Thomas et al., 2012).

### Terminal repeat regions

We initially noticed repeat regions at some of the chromosome ends. We used the Tandem Repeats Finder (v. 4.09) to identify terminal repeat regions (Benson, 1999). We scanned 24 kb of each chromosome end independently for repeats. For many of the chromosome scaffolds this yielded the same repeat in the 3’ direction. Inspection of the S1 region led to the discovery of the similar repeat, albeit more degenerate. Thereafter, we also scanned the entire scaffolds for the conserved repeat. However, only S1 contained the 16mer repeat at its center.

### Predicted Secreted Proteins (PSPs) filtering

We used SignalP v.6.0h on slow sequential and DeepTMHMM v.1.0.42 to predict the signal peptides and transmembrane domains in the protein coding genes respectively (Hallgren et al., 2022; Teufel et al., 2022). We used custom R scripts employed in R v.4.4.1 and RStudio build 421 and python scripts employed in python v.3.10.8 to filter out the genes with Signal Peptide and without any transmembrane domains. The custom R and python scripts can be obtained in this github repository.

### Ortholog Analysis

Orthofinder v. 2.5.4 was used to find orthologs of all annotated proteins in *M. hapla* against 72 nematode species with Tardigrade as outgroup (Emms & Kelly, 2019). We used the option - msa to produce multiple sequence alignments and phylogenetic trees for all OrthoGroups. We started from a previous OrthoFinder study (Mota et al., 2023) and we replaced the former *M. hapla* set of predicted proteins by this new one.

### Functional annotation of CAZymes

For functional annotation of CAZymes (GH, PL and GT), we used dbCAN3 using DIAMOND, HMMER via CAZY and dbCAN-sub (Zheng et al., 2023) and we further verified the annotations with InterproScan and EggNOG mapper. Carbohydrate Esterases (CEs) were not annotated by dbCAN3, so we used the predictions from BLAST2GO, InterproScan and EggNOG-mapper to annotate these proteins.

### Horizontal gene transfer (HGT) analysis

For HGT analysis, we used the AvP (Alienness vs Predictor) pipeline with a database constructed from UNIREF90 to select, extract and detect the HGT candidates among the predicted secreted proteins (Koutsovoulos et al., 2022).

### Phylogenetic trees for CAZymes

For construction of phylogenetic trees, we first used Clustal Omega for the multiple sequence alignment of each CAZyme family with their respective outgroups. Then we used Iqtree to construct the Maximum Likelihood tree under the best-fit model with ultrafast bootstrap that generated 1000 pseudo-replicates to assess support for every internal node. The command used was: iqtree -s input_alignment.phylip -B 1000. Finally, the tree was uploaded to iTOL to make further customizations.

### Hypergeometric Test

This analysis was done using R (v4.2.2) using tidyverse (v1.3.2) and purr (0.3.4) packages (Github link). The chromosomal domain content (“HRZ” and “LRZ”) was based on the genetic linkage map (**Supp. Table 2**). Gene start and end positions, obtained from genome annotation, were matched to chromosomal domains by ensuring that a gene’s entire length falls within a single domain interval. Genes overlapping multiple domains were assigned to each matching domain, and genes not directly overlapping any domain were assigned to the nearest domain based on the minimum distance between gene boundaries and domain boundaries. We then calculated the total number of genes in each domain (N_domain) and total number genes (N_total). After this, we extracted PSPs per domain (k_domain) and total number of PSPs genomewide (k_total). We followed this with an enrichment test, where we tested enrichment of PSPs in each domain using a one-tailed hypergeometric test (R’s *phyper* function), with the null hypothesis that PSPs are randomly distributed across domains. For a give domain (LRZ/HRZ), the test conducted using significance threshold of α=0.05

*P(X≥k_domain) = phyper(k_domain -1, N_domain, N_total - N_domain, k_total,lower.tail = FALSE)*,

where, N_domain = number of genes in domain

k_domain = number of PSPs in domain

N_total = total number of genes

k_total = total number of PSPs. Same process was repeated for *C. elegans* orthologs.

### Mitochondrial genome assembly and annotation

In the draft assembly, Contig 35l, with a coverage of 1750x was identified as mitochondria. To assemble a linear mitochondrial genome, we used MitoZ v.3.6 with Illumina DNA sequences (Meng et al., 2019). We used Mitoz all commands with clade option nematoda and genetic code 5. This was followed by annotation using the command Mitoz annotate. Although this process gave us the core genes in the mitochondrial genome, it failed to annotate most of the tRNA genes. For this, we used MITOS2 from web based Galaxy v.24.14 to reannotate the linear mitochondrial genome (Bernt et al., 2013). MITOS2 was able to annotate some of the missing tRNAs, but we only obtained 10 out of 22tRNAs using this method. We then downloaded the mitochondrial tRNA genes from *M. graminicola* from GenBank accession number KJ139963.1. We blasted these tRNA sequences against the linear mitochondrial genome of *M. hapla* and obtained 17 out of 22 tRNA sequences. We then employed Circos to plot the diagram for the mitochondrial genome.

### DNA Fluorescent in-situ hybridization (FISH) of the candidate terminal repeat

*M. hapla* genomic DNA was isolated from 50 collected egg sacs using Qiagen Blood and Tissue kit (Qiagen) according to the manufacturer’s procedure with slight modifications. Tissue was homogenized for 30 sec using electric homogenizer and pestle and incubated O/N at 56 °C. Sample was then treated with RNase A for 5 min at RT and final elution was in 100 µL of the elution buffer. For PCR procedures obtained DNA was diluted to concentration of 0,02 ng/µL. To produce probes corresponding to the 16 bp repeat (GTTTAAAAGGCCCAAG) identified at scaffold ends, we extended primers to encompass more than one monomer to avoid primer dimers as much as possible (Mhap_tel_4R AAGATTTAAAAGGCCCAAGATTTAAAAGAC Mhap_tel_4L CTTTTAAACCTTGGGTCTTTTAAACCTTG). FISH probes were prepared according to the previously described procedure (Despot-Slade et al., 2021) with 61.7 °C as annealing temperature. Prepared probes were cleaned using QIAquick PCR Purification Kit (Qiagen) and eluted in 50 µL of nuclease-free water. They were tested on 1% agarose gel to check their concentration and length **(Supp. Fig. 11**).

Whole *M. hapla* females were squashed onto slides and different fixatives were tested to optimize chromosome morphology. Ethanol and acetic acid fixative proved to be optimal for both preservation of chromosome morphology and FISH analyses and slide preparation was done as described (Gržan et al., 2020). The FISH procedure was done according to established protocols (Despot-Slade et al., 2021; Mota et al., 2023) except for skipping the pretreatment step of specimens in 45 % acetic acid as here the slides were fixed in a mixture already containing acetic acid. Similarly, as *M. incognita*, it was not possible to count chromosome numbers exactly and prometaphases/metaphases are rare. Telomeric signals had different intensity from very large to very discrete and consequently it was hard to count the precise number of certain signals. Best evaluation was obtained on elongated chromosomes (prometaphase).

## Acknowledgements

We would like to thank Michael Winters and Dave Lunt for helpful discussions. We would also like to thank the University of California Davis farm server system and Wageningen University HPC cluster system for their support with the data analysis.

## Author contributions

Conceived and administered the project: SS, MGS, VMW; DNA extraction and sequencing: PS, VMW; Assembled and annotated the genome: PS, SvdR, ADP, DD; Conducted genetic analysis: MIM, MLV, MGS, VMW; Telomere repeat analysis: EGD, AZM; Wrote the draft manuscript: PS; Revised the manuscript: MGS, VW, SS.

## Data availability

All sequencing data have been deposited in the Sequence Read Archive (SRA) at the National Center for Biotechnology Information (NCBI). The HiFi, Nanopore, Illumina, HiC, and Iso-seq data were deposited under Bioproject PRJNA1265270. The processed genome and annotation data is available under Bioproject PRJNA1265270. The supplementary and source data is publicly available at Zenodo research repository (https://doi.org/10.5281/zenodo.15484846). The scripts used for generation of figures and data analysis can be found in: https://github.com/Siddique-Lab/Mhapla_genome_paper; https://git.wur.nl/stefan.vanderuitenbeek/rnaseq_variant_calling_snakemake_pipeline/ ; https://git.wur.nl/published_papers/vw9_genome

## References

Bali, S., & Gleason, C. (2024). Unveiling the Diversity: Plant Parasitic Nematode Effectors and Their Plant Interaction Partners. Molecular Plant-Microbe Interactions: MPMI, 37(3), 179–189.

Bali, S., Hu, S., Vining, K., Brown, C., Mojtahedi, H., Zhang, L., Gleason, C., & Sathuvalli, V. (2021). Nematode Genome Announcement: Draft Genome of *Meloidogyne chitwoodi*, an Economically Important Pest of Potato in the Pacific Northwest. Molecular Plant-Microbe Interactions: MPMI, 34(8), 981–986.

Beek, J. G. van der, Los, J. A., & Pijnacker, L. P. (1998). Cytology of parthenogenesis of five Meloidogyne species. Fundamental and Applied Nematology, 21(4), Article 4.

Benson, G. (1999). Tandem repeats finder: A program to analyze DNA sequences. Nucleic Acids Research, 27(2), 573–580.

Bernt, M., Donath, A., Jühling, F., Externbrink, F., Florentz, C., Fritzsch, G., Pütz, J., Middendorf, M., & Stadler, P. F. (2013). MITOS: Improved *de novo* metazoan mitochondrial genome annotation. Molecular Phylogenetics and Evolution, 69(2), 313–319.

Bird, D. McK., Williamson, V. M., Abad, P., McCarter, J., Danchin, E. G. J., Castagnone-Sereno, P., & Opperman, C. H. (2009). The Genomes of Root-Knot Nematodes. Annual Review of Phytopathology, 47(1), 333–351.

Blanc-Mathieu, R., Perfus-Barbeoch, L., Aury, J.-M., Rocha, M. D., Gouzy, J., Sallet, E., Martin-Jimenez, C., Bailly-Bechet, M., Castagnone-Sereno, P., Flot, J.-F., Kozlowski, D. K., Cazareth, J., Couloux, A., Silva, C. D., Guy, J., Kim-Jo, Y.-J., Rancurel, C., Schiex, T., Abad, P., … Danchin, E. G. J. (2017). Hybridization and polyploidy enable genomic plasticity without sex in the most devastating plant-parasitic nematodes. PLOS Genetics, 13(6), e1006777.

Buchfink, B., Reuter, K., & Drost, H.-G. (2021). Sensitive protein alignments at tree-of-life scale using DIAMOND. Nature Methods, 18(4), 366–368.

Burga, A., Ben-David, E., Lemus Vergara, T., Boocock, J., & Kruglyak, L. (2019). Fast genetic mapping of complex traits in C. elegans using millions of individuals in bulk. Nature Communications, 10(1), 2680.

Cantalapiedra, C. P., Hernández-Plaza, A., Letunic, I., Bork, P., & Huerta-Cepas, J. (2021). egg- NOG-mapper v2: Functional Annotation, Orthology Assignments, and Domain Prediction at the Metagenomic Scale. Molecular Biology and Evolution, 38(12), 5825–5829.

Challis, R., Richards, E., Rajan, J., Cochrane, G., & Blaxter, M. (2020). BlobToolKit – Interactive Quality Assessment of Genome Assemblies. G3 Genes|Genomes|Genetics, 10(4), 1361–1374.

Cheng, H., Concepcion, G. T., Feng, X., Zhang, H., & Li, H. (2021). Haplotype-resolved de novo assembly using phased assembly graphs with hifiasm. Nature Methods, 18(2), 170–175.

Croll, D., Lendenmann, M. H., Stewart, E., & McDonald, B. A. (2015). The Impact of Recombination Hotspots on Genome Evolution of a Fungal Plant Pathogen. Genetics, 201(3), 1213–1228.

Dai, D., Xie, C., Zhou, Y., Bo, D., Zhang, S., Mao, S., Liao, Y., Cui, S., Zhu, Z., Wang, X., Li, F., Peng, D., Zheng, J., & Sun, M. (2023). Unzipped chromosome-level genomes reveal allopolyploid nematode origin patterns as unreduced gamete hybridization. Nature Communications, 14(1), 7156.

Despot-Slade, E., Mravinac, B., Širca, S., Castagnone-Sereno, P., Plohl, M., & Meštrović, N. (2021). The Centromere Histone Is Conserved and Associated with Tandem Repeats Sharing a Conserved 19-bp Box in the Holocentromere of Meloidogyne Nematodes. Molecular Biology and Evolution, 38(5), 1943–1965.

Dudchenko, O., Batra, S. S., Omer, A. D., Nyquist, S. K., Hoeger, M., Durand, N. C., Shamim, M. S., Machol, I., Lander, E. S., Aiden, A. P., & Aiden, E. L. (2017). De novo assembly of the Aedes aegypti genome using Hi-C yields chromosome-length scaffolds. Science, 356(6333), 92–95.

Durand, N. C., Shamim, M. S., Machol, I., Rao, S. S. P., Huntley, M. H., Lander, E. S., & Aiden, E. L. (2016). Juicer provides a one-click system for analyzing loop-resolution Hi-C experiments. Cell Systems, 3(1), 95–98.

Emms, D. M., & Kelly, S. (2019). OrthoFinder: Phylogenetic orthology inference for comparative genomics. Genome Biology, 20(1), 238.

Escobar, C., Barcala, M., Cabrera, J., & Fenoll, C. (2015). Chapter One—Overview of Root-Knot Nematodes and Giant Cells. In C. Escobar & C. Fenoll (Eds.), Advances in Botanical Research (Vol. 73, pp. 1–32). Academic Press.

Flynn, J. M., Hubley, R., Goubert, C., Rosen, J., Clark, A. G., Feschotte, C., & Smit, A. F. (2020). RepeatModeler2 for automated genomic discovery of transposable element families. Proceedings of the National Academy of Sciences, 117(17), 9451–9457.

Fouché, S., Plissonneau, C., & Croll, D. (2018). The birth and death of effectors in rapidly evolving filamentous pathogen genomes. Current Opinion in Microbiology, 46, 34–42.

Gabriel, L., Brůna, T., Hoff, K. J., Ebel, M., Lomsadze, A., Borodovsky, M., & Stanke, M. (2024). BRAKER3: Fully automated genome annotation using RNA-seq and protein evidence with GeneMark-ETP, AUGUSTUS and TSEBRA. bioRxiv, 2023.06.10.544449.

Goldstein, P., & Triantaphyllou, A. C. (1978). Occurrence of synaptonemal complexes and recombination nodules in a meiotic race of Meloidogyne hapla and their absence in a mitotic race. Chromosoma, 68(1), 91–100.

Gonzalez de la Rosa, P. M., Thomson, M., Trivedi, U., Tracey, A., Tandonnet, S., & Blaxter, M. (2021). A telomere-to-telomere assembly of Oscheius tipulae and the evolution of rhabditid nematode chromosomes. G3 Genes|Genomes|Genetics, 11(1), jkaa020.

Grbić, M., Van Leeuwen, T., Clark, R. M., Rombauts, S., Rouzé, P., Grbić, V., Osborne, E. J., Dermauw, W., Thi Ngoc, P. C., Ortego, F., Hernández-Crespo, P., Diaz, I., Martinez, M., Navajas, M., Sucena, É., Magalhães, S., Nagy, L., Pace, R. M., Djuranović, S., … Van de Peer, Y. (2011). The genome of Tetranychus urticae reveals herbivorous pest adaptations. Nature, 479(7374), 487– 492.

Griffin, S. (2023). Sogriffin98/Contamination_of_Genomes_using_BlobToolKit [Computer software]. https://github.com/sogriffin98/Contamination_of_Genomes_using_BlobToolKit (Original work published 2023)

Gržan, T., Despot-Slade, E., Meštrović, N., Plohl, M., & Mravinac, B. (2020). CenH3 distribution reveals extended centromeres in the model beetle Tribolium castaneum. PLOS Genetics, 16(10), e1009115. 10.1371/journal.pgen.1009115

Guo, Y., Fudali, S., Gimeno, J., DiGennaro, P., Chang, S., Williamson, V. M., Bird, D. M., & Nielsen, D. M. (2017). Networks Underpinning Symbiosis Revealed Through Cross-Species eQTL Mapping. Genetics, 206(4), 2175–2184.

Haegeman, A., Jones, J. T., & Danchin, E. G. J. (2011). Horizontal Gene Transfer in Nematodes: A Catalyst for Plant Parasitism? Molecular Plant-Microbe Interactions®, 24(8), 879–887.

Hallgren, J., Tsirigos, K. D., Pedersen, M. D., Armenteros, J. J. A., Marcatili, P., Nielsen, H., Krogh, A., & Winther, O. (2022). *DeepTMHMM predicts alpha and beta transmembrane proteins using deep neural networks* (p. 2022.04.08.487609). bioRxiv.

Hillers, K. J., & Villeneuve, A. M. (2003). Chromosome-Wide Control of Meiotic Crossing over in C. elegans. Current Biology, 13(18), 1641–1647.

Hillier, L. W., Miller, R. D., Baird, S. E., Chinwalla, A., Fulton, L. A., Koboldt, D. C., & Waterston, R. H. (2007). Comparison of C. elegans and C. briggsae Genome Sequences Reveals Extensive Conservation of Chromosome Organization and Synteny. PLOS Biology, 5(7), e167.

Hinch, R., Donnelly, P., & Hinch, A. G. (2023). Meiotic DNA breaks drive multifaceted mutagenesis in the human germ line. Science, 382(6674), eadh2531.

Hugot, J.-P., Baujard, P., & Morand, S. (2001). Biodiversity in helminths and nematodes as a field of study: An overview. Nematology, 3(3), 199–208.

Jagdale, S., Rao, U., & Giri, A. P. (2021). Effectors of Root-Knot Nematodes: An Arsenal for Successful Parasitism. Frontiers in Plant Science, 12.

Jones, J. T., Haegeman, A., Danchin, E. G. J., Gaur, H. S., Helder, J., Jones, M. G. K., Kikuchi, T., Manzanilla-López, R., Palomares-Rius, J. E., Wesemael, W. M. L., & Perry, R. N. (2013). Top 10 plant-parasitic nematodes in molecular plant pathology. Molecular Plant Pathology, 14(9), 946–961.

Jones, P., Binns, D., Chang, H.-Y., Fraser, M., Li, W., McAnulla, C., McWilliam, H., Maslen, J., Mitchell, A., Nuka, G., Pesseat, S., Quinn, A. F., Sangrador-Vegas, A., Scheremetjew, M., Yong, S.-Y., Lopez, R., & Hunter, S. (2014). InterProScan 5: Genome-scale protein function classification. Bioinformatics, 30(9), 1236–1240.

Killick, R., & Eckley, I. A. (2014). changepoint: An R Package for Changepoint Analysis. Journal of Statistical Software, 58, 1–19. 10.18637/jss.v058.i03

Kim, J., Yang, R., Chang, C., Park, Y., & Tucker, M. L. (2018). The root-knot nematode *Meloidogyne incognita* produces a functional mimic of the Arabidopsis INFLORESCENCE DEFICIENT IN ABSCISSION signaling peptide. Journal of Experimental Botany, 69(12), 3009–3021.

Koren, S., Walenz, B. P., Berlin, K., Miller, J. R., Bergman, N. H., & Phillippy, A. M. (2017). Canu: Scalable and accurate long-read assembly via adaptive k-mer weighting and repeat separation. Genome Research, 27(5), 722–736.

Koutsovoulos, G. D., Noriot, S. G., Bailly-Bechet, M., Danchin, E. G. J., & Rancurel, C. (2022). AvP: A software package for automatic phylogenetic detection of candidate horizontal gene transfers. PLOS Computational Biology, 18(11), e1010686.

Laurent, B., Palaiokostas, C., Spataro, C., Moinard, M., Zehraoui, E., Houston, R. D., & Foulongne-Oriol, M. (2017). High-resolution mapping of the recombination landscape of the phytopathogen Fusarium graminearum suggests two-speed genome evolution. Molecular Plant Pathology, 19(2), 341–354.

Li, H. (2013). *Aligning sequence reads, clone sequences and assembly contigs with BWA-MEM* (No. arXiv:1303.3997). arXiv.

Li, H. (2018). Minimap2: Pairwise alignment for nucleotide sequences. Bioinformatics, 34(18), 3094–3100.

Li, M., Zhu, C., Xu, Z., Xu, M., Kuang, Y., Hou, X., Huang, X., Lv, M., Liu, Y., Zhang, Y., Xu, Z., Han, X., Wang, S., Shi, Y., Guang, S., & Li, F. (2024). Structural basis for C. elegans pairing center DNA binding specificity by the ZIM/HIM-8 family proteins. Nature Communications, 15(1), 10355.

Lieberman-Aiden, E., van Berkum, N. L., Williams, L., Imakaev, M., Ragoczy, T., Telling, A., Amit, I., Lajoie, B. R., Sabo, P. J., Dorschner, M. O., Sandstrom, R., Bernstein, B., Bender, M. A., Groudine, M., Gnirke, A., Stamatoyannopoulos, J., Mirny, L. A., Lander, E. S., & Dekker, J. (2009). Comprehensive mapping of long-range interactions reveals folding principles of the human genome. *Science (New York*, N.Y.), 326(5950), 289–293. 10.1126/science.1181369

Lin, C.-J., & Siddique, S. (2024). Parasitic nematodes: Dietary habits and their implications. Trends in Parasitology, 40(3), 230–240.

Liu, Q. L., Thomas, V. P., & Williamson, V. M. (2007a). Meiotic Parthenogenesis in a Root-Knot Nematode Results in Rapid Genomic Homozygosity. Genetics, 176(3), 1483–1490.

Liu, Q. L., Thomas, V. P., & Williamson, V. M. (2007b). Meiotic Parthenogenesis in a Root-Knot Nematode Results in Rapid Genomic Homozygosity. Genetics, 176(3), 1483–1490.

Liu, Q. L., & Williamson, V. M. (2006). Host-Specific Pathogenicity and Genome Differences between Inbred Strains of Meloidogyne hapla. Journal of Nematology, 38(1), 158–164.

Manni, M., Berkeley, M. R., Seppey, M., & Zdobnov, E. M. (2021). BUSCO: Assessing Genomic Data Quality and Beyond. Current Protocols, 1(12), e323. 10.1002/cpz1.323

Marçais, G., & Kingsford, C. (2011). A fast, lock-free approach for efficient parallel counting of occurrences of k-mers. Bioinformatics, 27(6), 764–770.

Melakeberhan, H., Douches, D., & Wang, W. (2012). Interactions of Selected Potato Cultivars and Populations of Meloidogyne hapla Adapted to the Midwest U.S. Soils. Crop Science, 52(3), 1132– 1137.

Meng, G., Li, Y., Yang, C., & Liu, S. (2019). MitoZ: A toolkit for animal mitochondrial genome assembly, annotation and visualization. Nucleic Acids Research, 47(11), e63.

Miller, J. R., Delcher, A. L., Koren, S., Venter, E., Walenz, B. P., Brownley, A., Johnson, J., Li, K., Mobarry, C., & Sutton, G. (2008). Aggressive assembly of pyrosequencing reads with mates. Bioinformatics, 24(24), 2818–2824.

Mishra, S., Hu, W., & DiGennaro, P. (2023). Root-Knot-Nematode-Encoded CEPs Increase Nitrogen Assimilation. Life, 13(10), 2020. 10.3390/life13102020

Mitkowski, N. A., & Abawi, G. S. (2003). Root-knot Nematode. The Plant Health Instructor, 3.

Molloy, B., Shin, D. S., Long, J., Pellegrin, C., Senatori, B., Vieira, P., Thorpe, P. J., Damm, A., Ahmad, M., Vermeulen, K., Derevnina, L., Wei, S., Sperling, A., Reyes Estévez, E., Bruty, S., de Souza, V. H. M., Kranse, O. P., Maier, T., Baum, T., & Eves-van den Akker, S. (2024). The origin, deployment, and evolution of a plant-parasitic nematode effectorome. PLoS Pathogens, 20(7), e1012395.

Mota, A.-P. Z., Koutsovoulos, G. D., Perfus-Barbeoch, L., Despot-Slade, E., Labadie, K., Aury, J.-M., Robbe-Sermesant, K., Bailly-Bechet, M., Belser, C., Péré, A., Rancurel, C., Kozlowski, D. K., Hassanaly-Goulamhoussen, R., Rocha, M. D., Noël, B., Mestrovic-Radan, N., Wincker, P., & Danchin, E. G. (2023). *Unzipped assemblies of polyploid root-knot nematode genomes reveal new kinds of unilateral composite telomeric repeats* (p. 2023.03.29.534350). bioRxiv.

Müller, M. C., Praz, C. R., Sotiropoulos, A. G., Menardo, F., Kunz, L., Schudel, S., Oberhänsli, S., Poretti, M., Wehrli, A., Bourras, S., Keller, B., & Wicker, T. (2019). A chromosome-scale genome assembly reveals a highly dynamic effector repertoire of wheat powdery mildew. The New Phytologist, 221(4), 2176–2189.

Opperman, C. H., Bird, D. M., Williamson, V. M., Rokhsar, D. S., Burke, M., Cohn, J., Cromer, J., Diener, S., Gajan, J., Graham, S., Houfek, T. D., Liu, Q., Mitros, T., Schaff, J., Schaffer, R., Scholl, E., Sosinski, B. R., Thomas, V. P., & Windham, E. (2008). Sequence and genetic map of Meloidogyne hapla: A compact nematode genome for plant parasitism. Proceedings of the National Academy of Sciences of the United States of America, 105(39), 14802–14807.

Páez, A. R. (2023). Plant-Parasitic Nematodes and Their Management: A Focus on New Nematicides. In *Nematodes—Ecology*, Adaptation and Parasitism. IntechOpen.

Prabh, N., & Rödelsperger, C. (2022). Multiple Pristionchus pacificus genomes reveal distinct evolutionary dynamics between de novo candidates and duplicated genes. Genome Research, 32(7), 1315–1327.

Quinlan, A. R., & Hall, I. M. (2010). BEDTools: A flexible suite of utilities for comparing genomic features. Bioinformatics, 26(6), 841–842.

Ranallo-Benavidez, T. R., Jaron, K. S., & Schatz, M. C. (2020). GenomeScope 2.0 and Smudgeplot for reference-free profiling of polyploid genomes. Nature Communications, 11(1), 1432.

Rehman, S., Gupta, V. K., & Goyal, A. K. (2016). Identification and functional analysis of secreted effectors from phytoparasitic nematodes. BMC Microbiology, 16(1), 48.

Ren, Y., Tseng, E., Smith, T. P. L., Hiendleder, S., Williams, J. L., & Low, W. Y. (2023). Long read isoform sequencing reveals hidden transcriptional complexity between cattle subspecies. BMC Genomics, 24(1), 108. 10.1186/s12864-023-09212-9

Rhie, A., Walenz, B. P., Koren, S., & Phillippy, A. M. (2020). Merqury: Reference-free quality, completeness, and phasing assessment for genome assemblies. Genome Biology, 21(1), 245.

Rockman, M. V., & Kruglyak, L. (2009). Recombinational Landscape and Population Genomics of Caenorhabditis elegans. PLOS Genetics, 5(3), e1000419.

Rödelsperger, C. (2024). Comparative Genomics of Sex, Chromosomes, and Sex Chromosomes in Caenorhabditis elegans and Other Nematodes. Methods in Molecular Biology (Clifton, N.J.), 2802, 455–472.

Ross, J. A., Koboldt, D. C., Staisch, J. E., Chamberlin, H. M., Gupta, B. P., Miller, R. D., Baird, S. E., & Haag, E. S. (2011). Caenorhabditis briggsae recombinant inbred line genotypes reveal inter-strain incompatibility and the evolution of recombination. PLoS Genetics, 7(7), e1002174.

Rutter, W. B., Franco, J., & Gleason, C. (2022). Rooting Out the Mechanisms of Root-Knot Nematode-Plant Interactions. Annual Review of Phytopathology, 60, 43–76.

Seidel, H. S., Rockman, M. V., & Kruglyak, L. (2008). Widespread Genetic Incompatibility in C. Elegans Maintained by Balancing Selection. Science, 319(5863), 589–594.

Siddique, S., Radakovic, Z. S., Hiltl, C., Pellegrin, C., Baum, T. J., Beasley, H., Bent, A. F., Chitambo, O., Chopra, D., Danchin, E. G. J., Grenier, E., Habash, S. S., Hasan, M. S., Helder, J., Hewezi, T., Holbein, J., Holterman, M., Janakowski, S., Koutsovoulos, G. D., … Eves-van den Akker, S. (2022). The genome and lifestage-specific transcriptomes of a plant-parasitic nematode and its host reveal susceptibility genes involved in trans-kingdom synthesis of vitamin B5. Nature Communications, 13(1), 6190.

Smit, A. F. A., Hubley, R., & Green, P. (2021). 2013*–2015*. *RepeatMasker Open-4.0.*

Spieth, J., Lawson, D., Davis, P., Williams, G., & Howe, K. (2018). Overview of gene structure in C. elegans. In WormBook: The Online Review of C. elegans Biology [Internet]. WormBook.

Szitenberg, A., Salazar-Jaramillo, L., Blok, V. C., Laetsch, D. R., Joseph, S., Williamson, V. M., Blaxter, M. L., & Lunt, D. H. (2017). Comparative Genomics of Apomictic Root-Knot Nematodes: Hybridization, Ploidy, and Dynamic Genome Change. Genome Biology and Evolution, 9(10), 2844–2861.

Tandonnet, S., Koutsovoulos, G. D., Adams, S., Cloarec, D., Parihar, M., Blaxter, M. L., & Pires-daSilva, A. (2019). Chromosome-Wide Evolution and Sex Determination in the Three-Sexed Nematode Auanema rhodensis. G3 Genes|Genomes|Genetics, 9(4), 1211–1230.

Teufel, F., Almagro Armenteros, J. J., Johansen, A. R., Gíslason, M. H., Pihl, S. I., Tsirigos, K. D., Winther, O., Brunak, S., von Heijne, G., & Nielsen, H. (2022). SignalP 6.0 predicts all five types of signal peptides using protein language models. Nature Biotechnology, 40(7), 1023–1025.

Thomas, V. P., Fudali, S. L., Schaff, J. E., Liu, Q., Scholl, E. H., Opperman, C. H., Bird, D. M., & Williamson, V. M. (2012). A Sequence-Anchored Linkage Map of the Plant–Parasitic Nematode Meloidogyne hapla Reveals Exceptionally High Genome-Wide Recombination. G3: Genes|Genomes|Genetics, 2(7), 815–824.

Thomas, V. P., & Williamson, V. M. (2013). Segregation and mapping in the root-knot nematode Meloidogyne hapla of quantitatively inherited traits affecting parasitism. Phytopathology, 103(9), 935–940.

Triantaphyllou, A. C. (1966). Polyploidy and reproductive patterns in the root-knot nematode Meloidogyne hapla. Journal of Morphology, 118(3), 403–413.

Triantaphyllou, A. C., & Hirschmann, H. (1980). Cytogenetics and Morphology in Relation to Evolution and Speciation of Plant-Parasitic Nematodes. Annual Review of Phytopathology, 18(Volume 18, 1980), 333–359.

Trudgill, D. L., & Blok, V. C. (2001). APOMICTIC, POLYPHAGOUS ROOT-KNOT NEMATODES: Exceptionally Successful and Damaging Biotrophic Root Pathogens. Annual Review of Phytopathology, 39(Volume 39, 2001), 53–77.

Verhoeven, A., Finkers-Tomczak, A., Prins, P., Valkenburg-van Raaij, D. R., van Schaik, C. C., Overmars, H., van Steenbrugge, J. J. M., Tacken, W., Varossieau, K., Slootweg, E. J., Kappers, I. F., Quentin, M., Goverse, A., Sterken, M. G., & Smant, G. (2023). The root-knot nematode effector MiMSP32 targets host 12-oxophytodienoate reductase 2 to regulate plant susceptibility. New Phytologist, 237(6), 2360–2374.

Walker, B. J., Abeel, T., Shea, T., Priest, M., Abouelliel, A., Sakthikumar, S., Cuomo, C. A., Zeng, Q., Wortman, J., Young, S. K., & Earl, A. M. (2014). Pilon: An Integrated Tool for Comprehensive Microbial Variant Detection and Genome Assembly Improvement. PLOS ONE, 9(11), e112963.

Wang, C., Lower, S., Thomas, V. P., & Williamson, V. M. (2010). Root-Knot Nematodes Exhibit Strain-Specific Clumping Behavior That Is Inherited as a Simple Genetic Trait. PLoS ONE, 5(12), e15148.

Winter, M. R., Taranto, A. P., Yimer, H. Z., Blundell, A. C., Siddique, S., Williamson, V. M., & Lunt, D. H. (2024). Phased chromosome-scale genome assembly of an asexual, allopolyploid root-knot nematode reveals complex subgenomic structure. PLOS ONE, 19(6), e0302506.

Yimer, H. Z., Luu, D. D., Coomer Blundell, A., Ercoli, M. F., Vieira, P., Williamson, V. M., Ronald, P. C., & Siddique, S. (2023). Root-knot nematodes produce functional mimics of tyrosine-sulfated plant peptides. Proceedings of the National Academy of Sciences, 120(29), e2304612120.

Yin, Y., Mao, X., Yang, J., Chen, X., Mao, F., & Xu, Y. (2012). dbCAN: A web resource for automated carbohydrate-active enzyme annotation. Nucleic Acids Research, 40(W1), W445– W451.

Yu, Z., Kim, Y., & Dernburg, A. F. (2016). Meiotic recombination and the crossover assurance checkpoint in *Caenorhabditis elegans*. Seminars in Cell & Developmental Biology, 54, 106–116.

Zdraljevic, S., Walter-McNeill, L., Bruni, G., Bloom, J. S., Leighton, D. H. W., Marquez, H., & Kruglyak, L. (2024). *A maternal-effect toxin-antidote element causes larval arrest in C. elegans* (p. 2024.04.26.591160). bioRxiv.

Zhang, X., Peng, H., Zhu, S., Xing, J., Li, X., Zhu, Z., Zheng, J., Wang, L., Wang, B., Chen, J., Ming, Z., Yao, K., Jian, J., Luan, S., Coleman-Derr, D., Liao, H., Peng, Y., Peng, D., & Yu, F. (2020). Nematode-Encoded RALF Peptide Mimics Facilitate Parasitism of Plants through the FERONIA Receptor Kinase. Molecular Plant, 13(10), 1434–1454.

Zheng, J., Ge, Q., Yan, Y., Zhang, X., Huang, L., & Yin, Y. (2023). dbCAN3: Automated carbohydrate-active enzyme and substrate annotation. Nucleic Acids Research, 51(W1), W115–W121.

