## Supplementary Figures for "A Chromosome-scale Genome Assembly of *Meloidogyne hapla* Reveals Localized Recombination Hotspots Enriched with Effector Proteins"

**List of Supplementary Figures**

| **Supp. Fig. 1** | **Blobplot/Contamination plot of draft assembly** |
| --- | --- |
| **Supp. Fig. 2** | **Mitochondrial genome of Meloidogyne hapla strain VW9** |
| **Supp. Fig. 3** | **Genome Profile of *Meloidogyne hapla* strain VW9** |
| **Supp. Fig. 4** | **HiC quality check and HiC assembly with ONT reads** |
| **Supp. Fig. 5** | **Genetic correlation between and within scaffolds** |
| **Supp Fig. 6** | **Distribution of CAZymes in the genome of *M. hapla*** |
| **Supp. Fig. 7** | **Phylogenetic tree and distribution of GH proteins** |
| **Supp. Fig. 8** | **Phylogenetic trees and distribution of CE and PL proteins** |
| **Supp. Fig. 9** | **Phylogenetic tree and distribution of GT proteins** |
| **Supp. Fig. 10** | **Distribution of genes across the genome of M. hapla** |
| **Supp. Fig. 11** | **Agarose gel image of prepared FISH probe** **with λ DNA for concentration assessment of Markers**. |

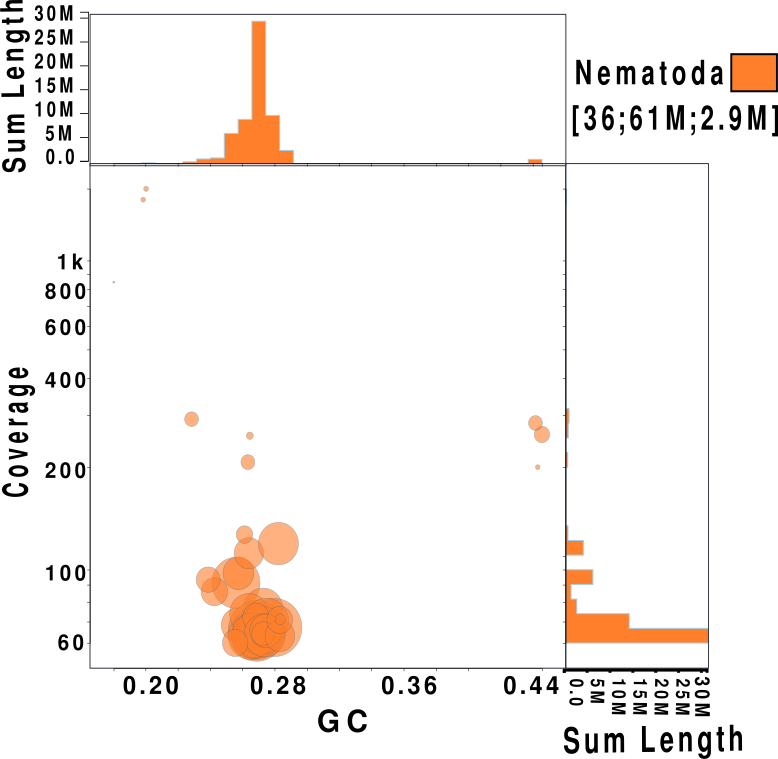

**Supp. Fig. 1:Blobplot/Contamination plot of draft assembly:** Each circle represents a contig sequence with size of the circle corresponding to contig size. The X-axis shows GC proportion and Y-axis shows base coverage. Circles are colored based on their taxonomic classification assigned via DIAMOND BLASTp against reference proteomes database and further refined using ‘best sum’ tax-rule. Histograms along the axes display the distribution of total assembly length. Contig 35 (with the highest coverage) corresponds to circular assembled mitochondrial DNA. The three contigs (Contig 27, Contig 33 and Contig 36), have high GC content and are repeat dense (~50%).

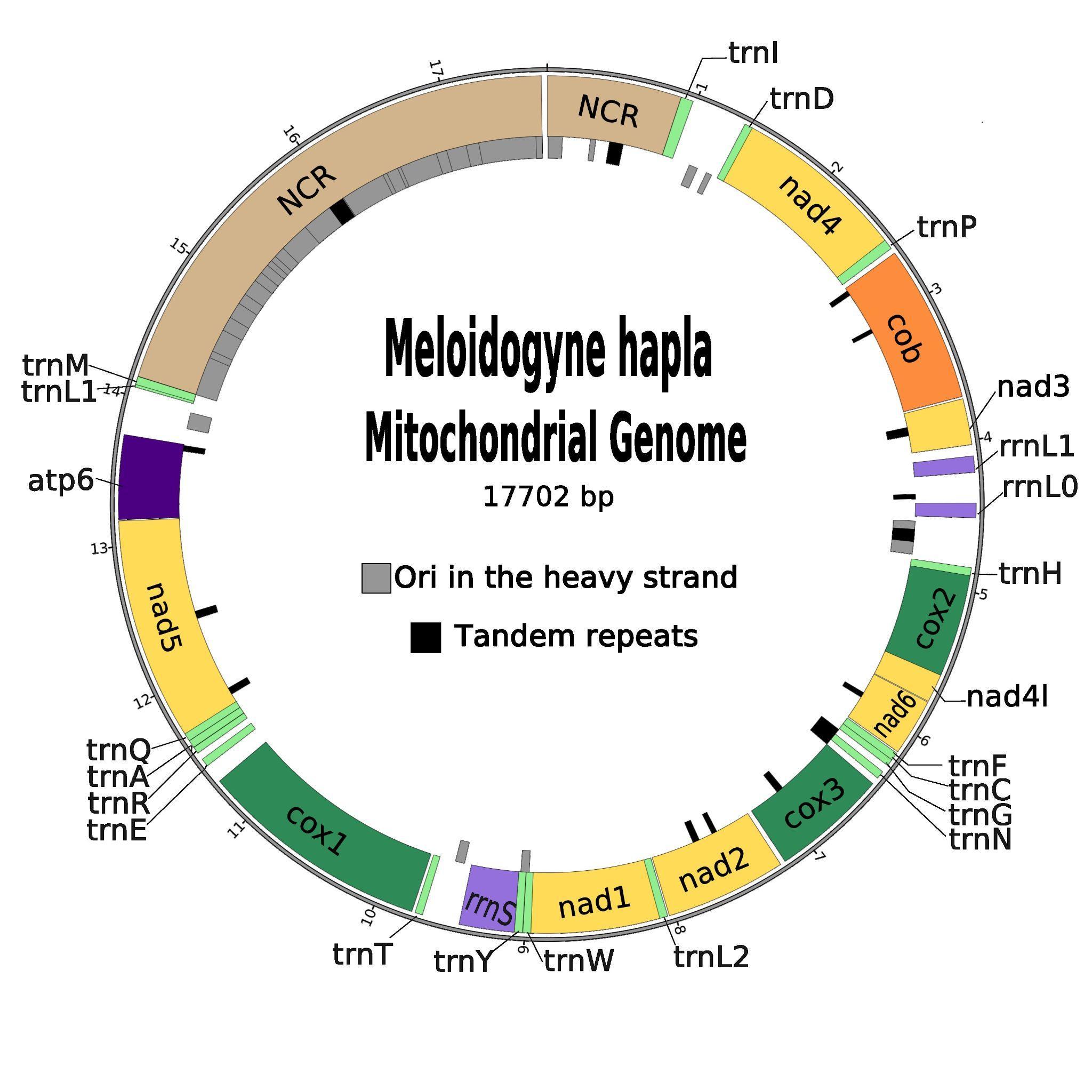

**Supp. Fig. 2. Mitochondrial genome of *Meloidogyne hapla* strain VW9:** The mitochondrial genome of *M. hapla* is 17.7 kb and AT rich with 12 protein coding genes, 17 tRNA genes and 2 large ribosomal subunit genes. NCR: Non Coding Region; NAD: Nicotinamide Adenine Dinucleotide; COX: Cytochrome c Oxidase; COB: Cytochrome B, NADL: Nicotinamide Adenine Dinucleotide Ligase, ATP(Adenosine Triphosphate).

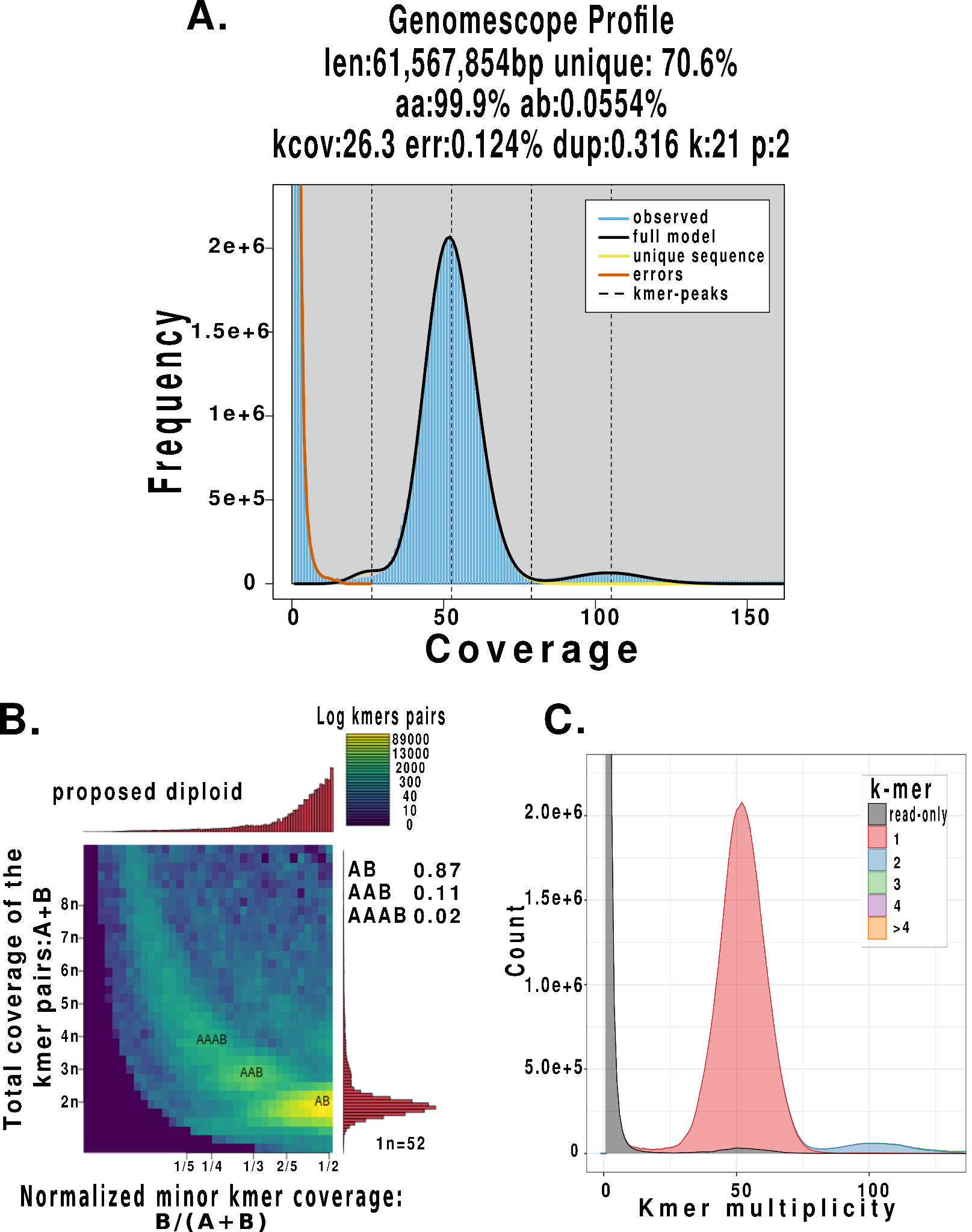

**Supp. Fig 3. Genome Profile of *Meloidogyne hapla* strain VW9: A.** Genomescope profile of HiFi Reads: The genome size of *M. hapla* is estimated to be 61.6 Mb. The single copy regions are at a coverage of approximately between 25 and 80, and the peak coverage position is at 52x. The plot shows that *M. hapla* strain VW9 is almost completely homozygous (99.9%). **B.** Smudgeplot profile produced with HiFi reads indicates that the genome is diploid. **C**. Merqury k-mer plot showing that almost all the information present in the HiFi reads has been captured in the genome assembly of *M. hapla.*

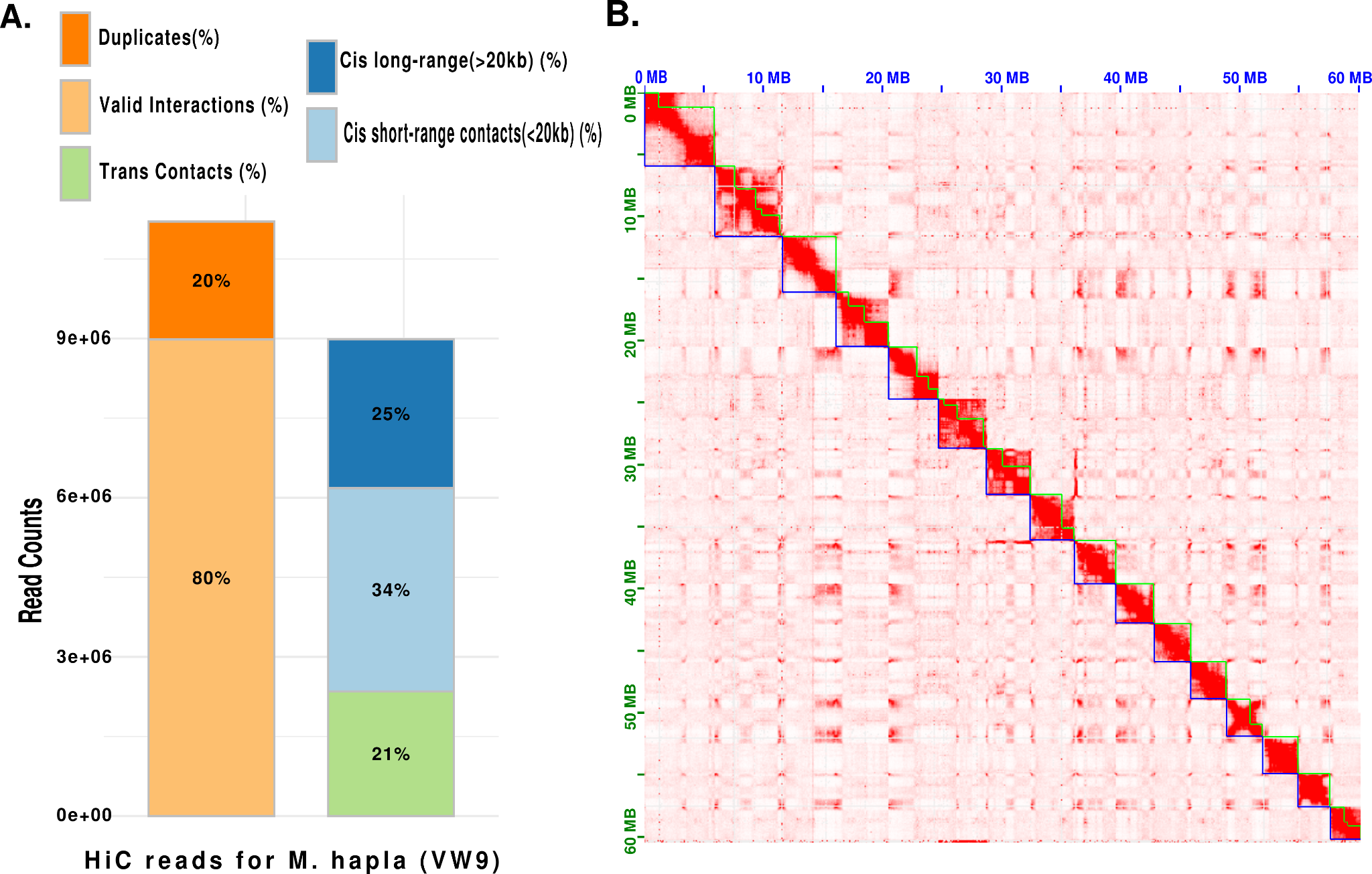

**Supp. Fig. 4. HiC quality check and HiC assembly with ONT reads: A.** Breakdown of valid pairs by duplicates and contact ranges. This plot characterizes the valid interaction pairs and highlights that the majority of interactions occur in the same chromosome, as is expected. The left bar shows distribution of valid interactions (80%) and duplicates (20%). The right bar shows the distribution of valid interactions by genomic distance: 25% are Cis long range contacts, 34% are cis short range contacts and 21% are trans contacts. **B.** HiC Contact Map of *M. hapla* made with Oxford Nanopore Reads using NECAT: The contact map shows 16 Scaffolds of *M. hapla* denoted by blue lines and 39 contigs denoted by green lines.

**Supp. Fig. 5: Genetic correlation between and within scaffolds:** Correlation plot among the 789 SNP markers used to generate genetic map. X- and y- axis represents the location of each marker. Diagonal boxes are the Chromosomes of M. hapla. Colour gradients represent the correlation between markers ranging from -1 (purple) to 1 (green).
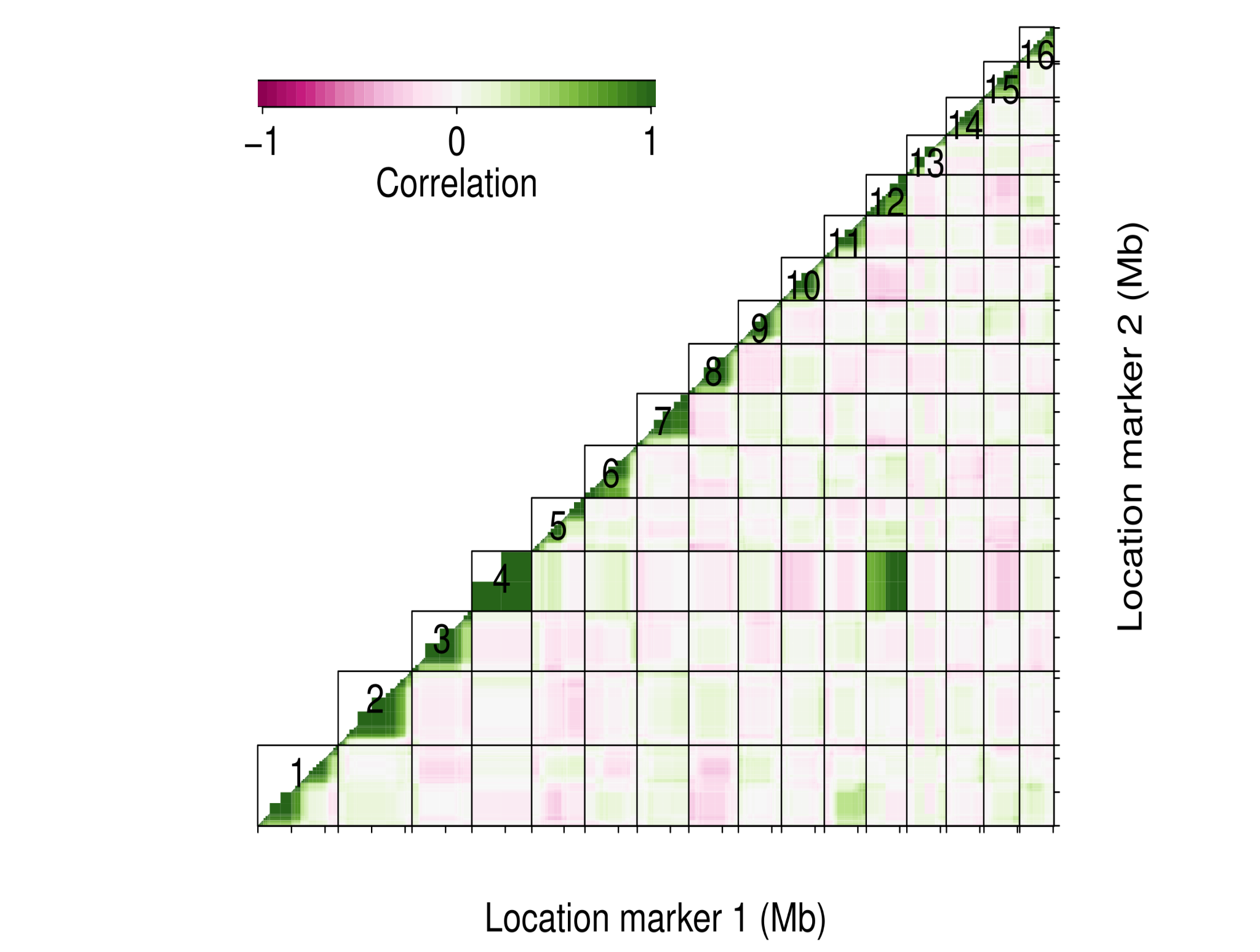

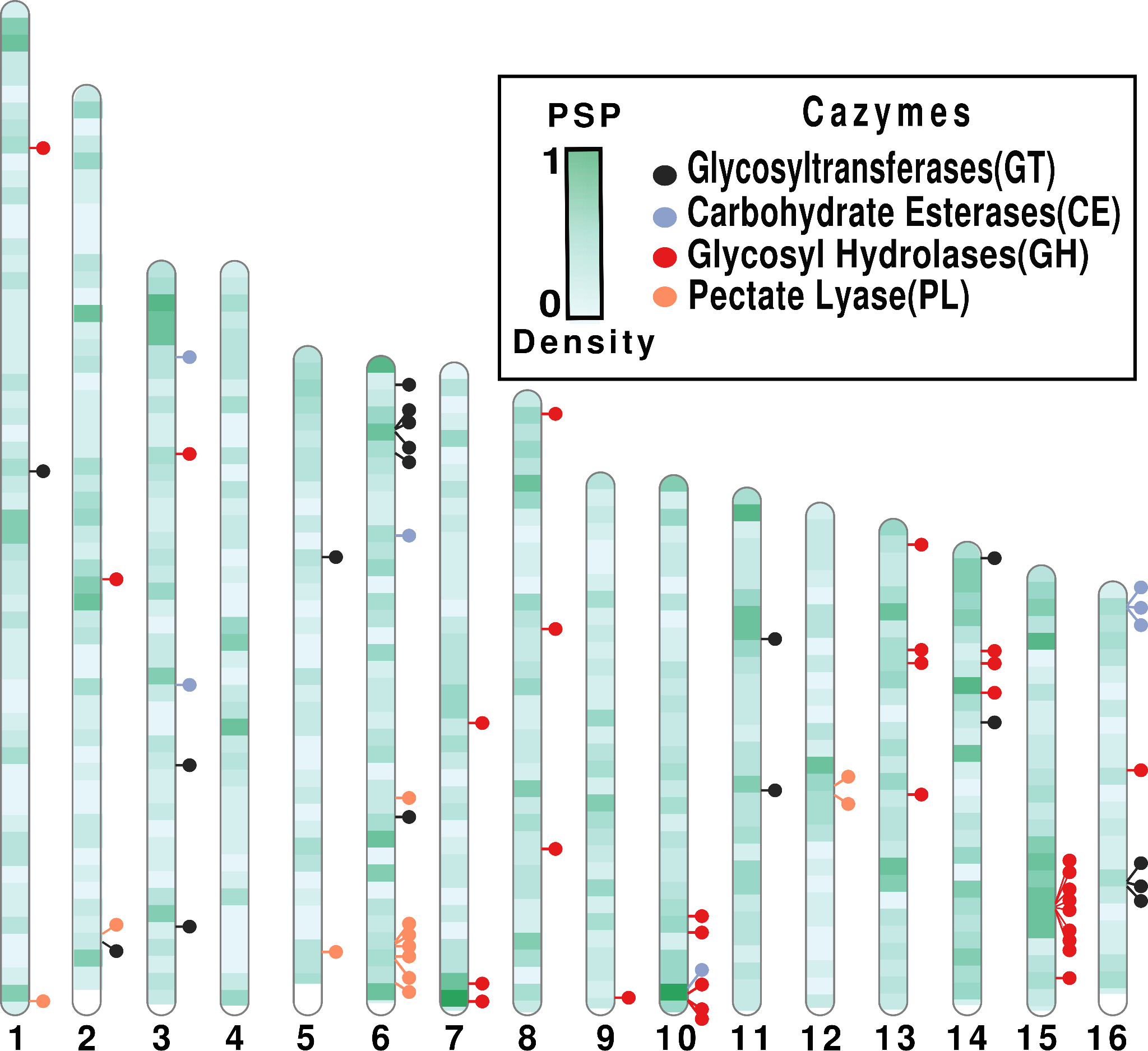

**Supp. Fig. 6. Distribution of secreted CAZymes in the genome of *M. hapla*:** Each chromosome is represented as a vertical bar and is divided into bins of 100KB. The PSP density (number of PSPs per 100 KB) is normalized to the values between 0 to 1. The positions of CAZymes within the genome are shown and are classified according to their family types: black for GT, light blue for CE, red for GH and orange for PL.

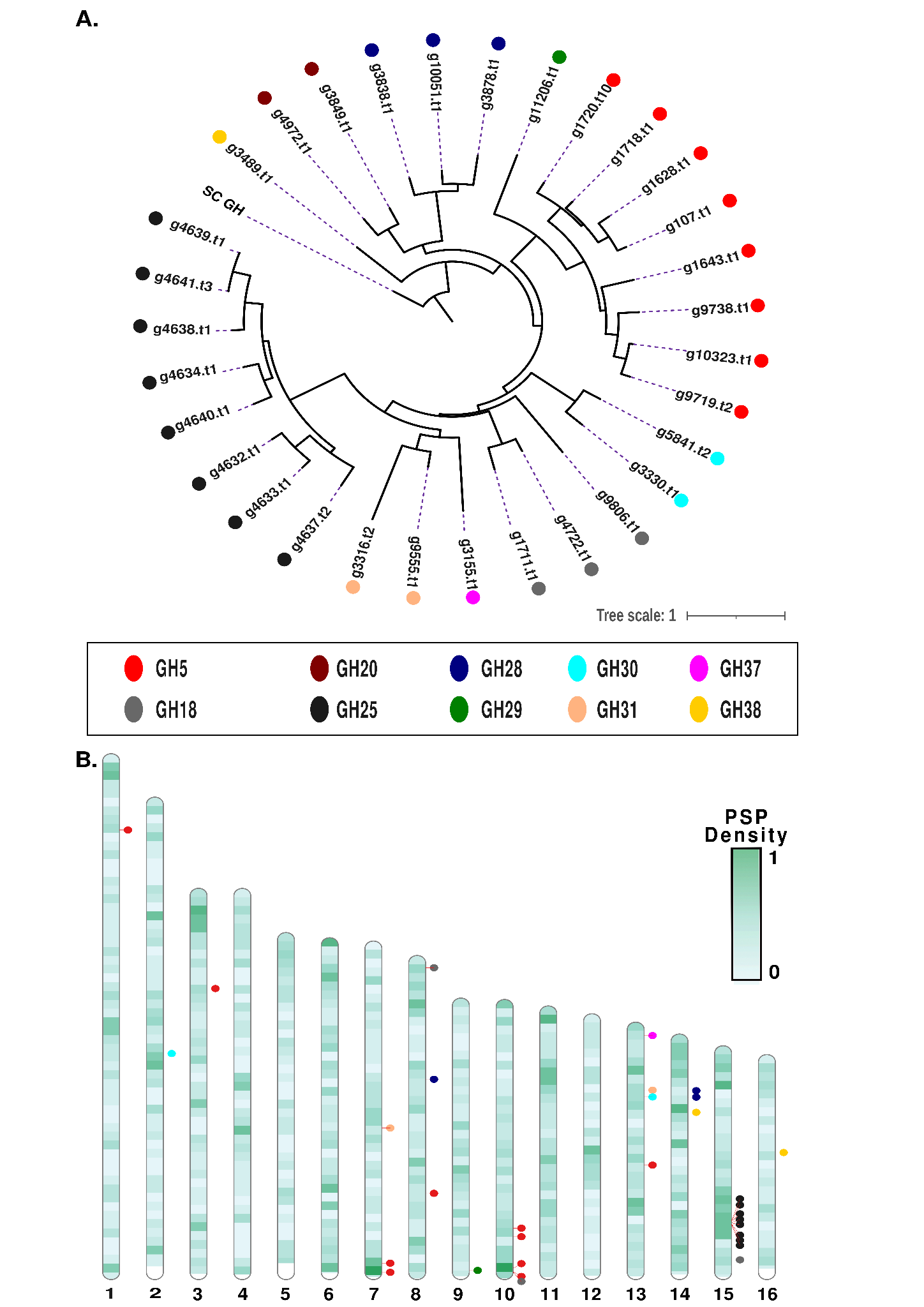

**Supp. Fig. 7: Phylogenetic tree and distribution of GH proteins: A.** Phylogenetic tree constructed by the maximum likelihood method shows the clustering of GH families. The outgroup is the GH11(Q9RI72) protein of *Streptomyces coelicolor.* Each color represents the subfamilies. Here, GH18 separates into two clusters as they belong to different sub families: g9806.t1 belongs to sub-family e374 and lacks catalytic annotation, while g4722.t1 and g1711.t1 belong to sub-families 2 and 4 respectively and are annotated as EC3.2.1.14 (chitinases) (**Supp. Table 10**) **B.** Position of GH families within the genome of *M. hapla.* GH proteins cluster based on their family classification. The physical proximity of different members of the same family within the scaffolds suggest possible gene cis-duplication events.

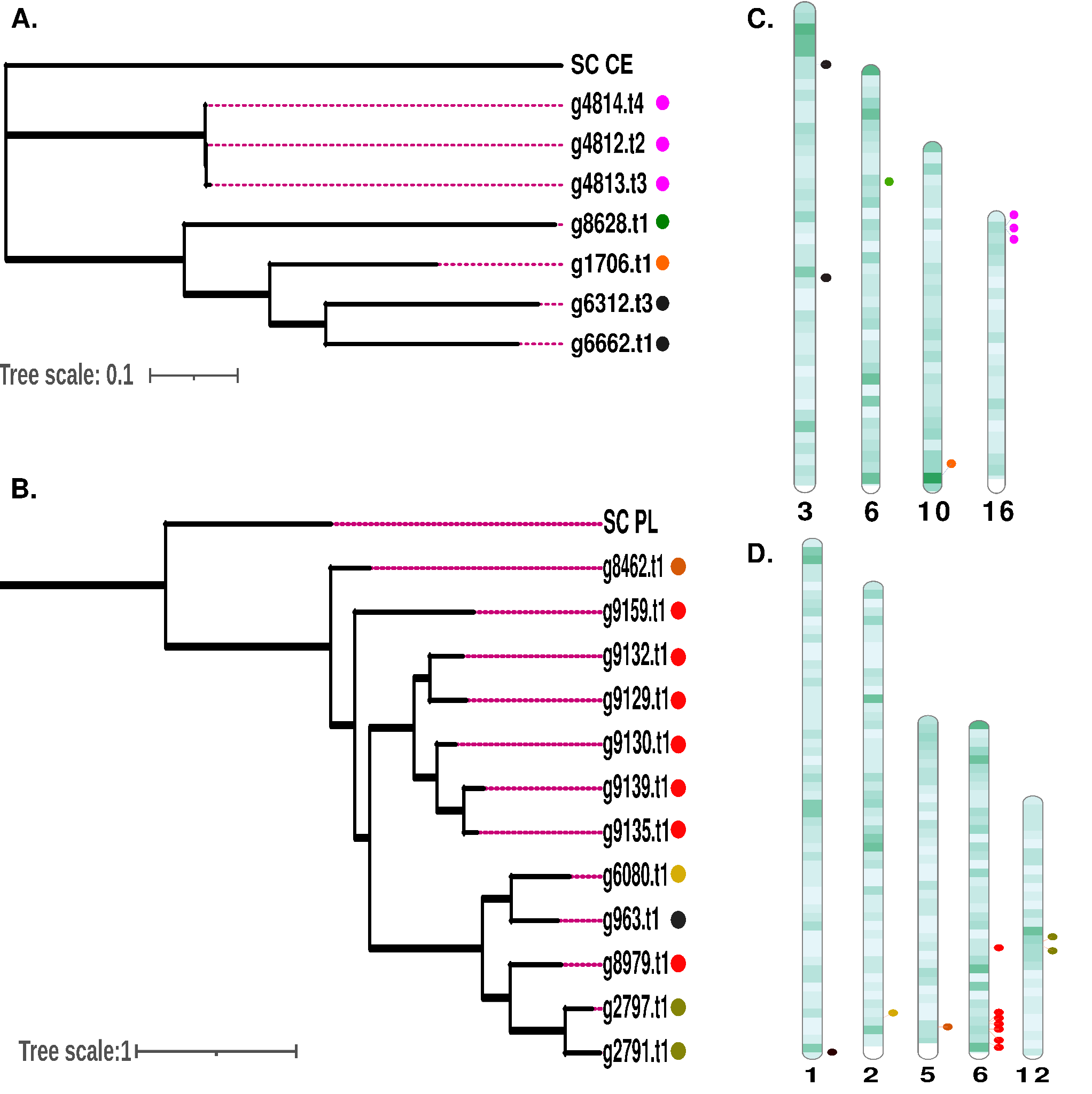

**Supp. Fig. 8. Phylogenetic trees and distribution of CE10 and PL5 proteins: A-B.** Phylogenetic tree constructed by the maximum likelihood method shows the clustering of CE and PL proteins. The outgroup is the respective proteins of *Streptomyces coelicolor:* Q93RU6 and Q9RKN7. Each color here represents the position of the protein in the chromosome. **C-D.** Positions of CE and PL proteins within the genome of *M. hapla.*

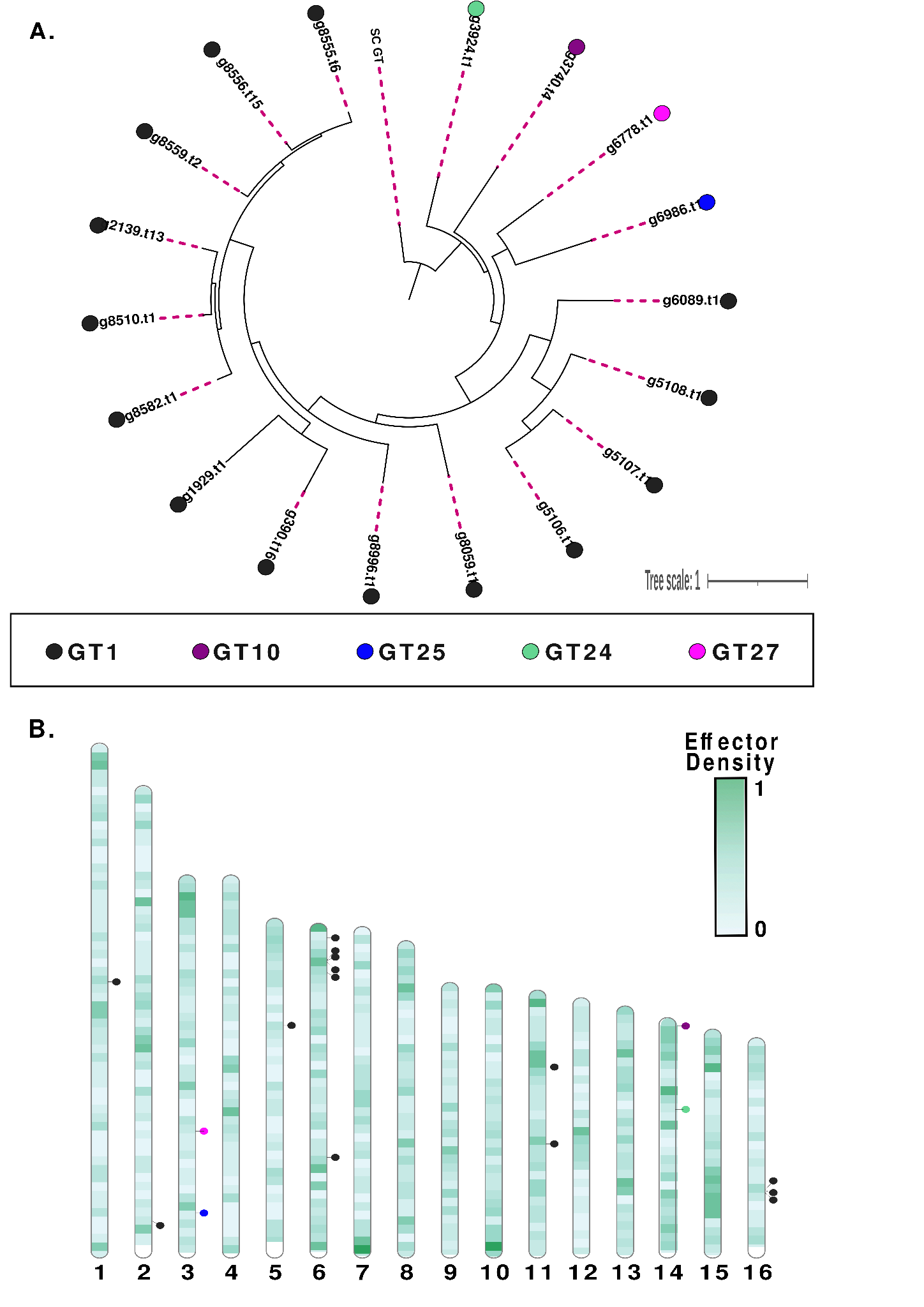

**Supp Fig. 9. Phylogenetic tree and distribution of GT proteins:** A. Phylogenetic tree constructed by the maximum likelihood method shows the clustering GT subfamilies. The outgroup is the GT1 (Q9KXK2) protein of *Streptomyces coelicolor.* Each color represents the families. **B.** Position of GT families within the genome of *M. hapla:* GT proteins cluster based on their family classification. The physical proximity of different members of the same family within the scaffolds suggest possible gene cis-duplication events.

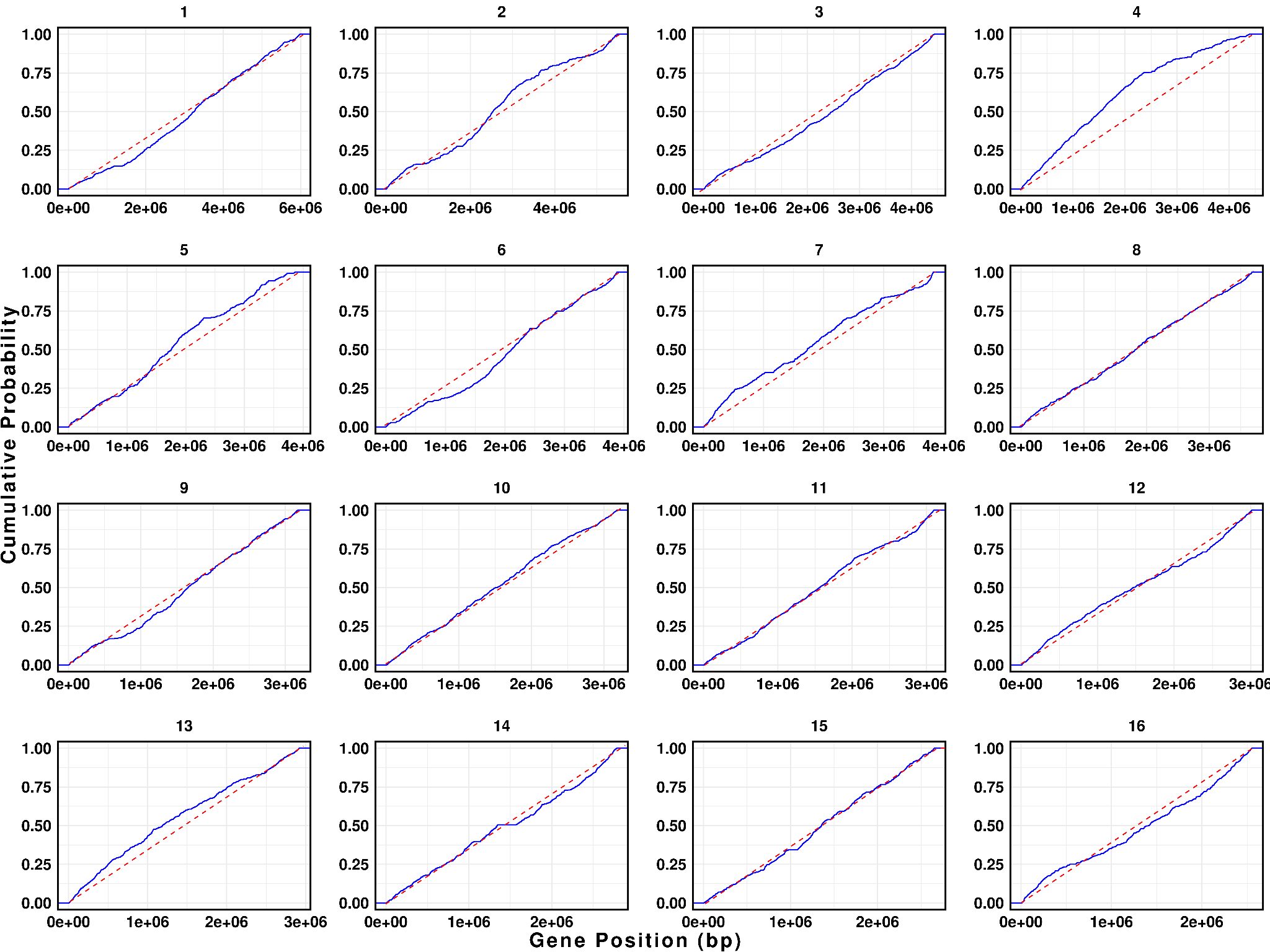

**Supp. Fig. 10.** **Distribution of genes across the genome of *M. hapla*.** The blue line represents the empirical cumulative distribution, showing the cumulative probability (or proportion) of genes positioned up to each point along the scaffold. The red dashed line indicates what the distribution would look like if genes were perfectly evenly distributed along the scaffold (theoretical uniform distribution). The X-axis displays the position along the scaffold in base pairs, reflecting the length of the scaffold, and the Y-axis represents the cumulative proportion of genes up to each position.

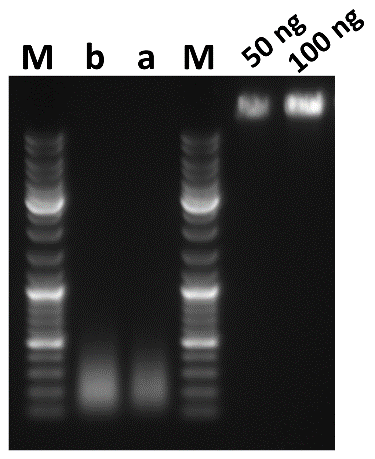

**Supp. Fig. 11**. **Agarose gel image of prepared FISH probe** **with λ DNA for concentration assessment of Markers**. M-marker, b-before PCR purification step, a-after PCR purification step.
